# Genome-wide screen in *Mycobacterium tuberculosis* infected macrophages reveals innate regulation of antibacterial mediators by IRF2

**DOI:** 10.1101/2025.09.26.678671

**Authors:** Allison W. Roberts, Linda N. Del Cid, Nicholas E. Garelis, Jeffery S. Cox

## Abstract

Controlling *Mycobacterium tuberculosis* (Mtb) infection requires a precisely balanced host inflammatory response. Too little inflammation leads to uncontrolled bacterial growth but exacerbated inflammation, activated by mediators such as TNF and type I IFN, inhibits effective antibacterial responses. How these immunopathological states are established is unknown. Deeper understanding of the pathways elicited upon initial Mtb infection of the host macrophage may reveal vital regulatory mechanisms that govern the subsequent inflammatory environment and ultimate resolution of infection. To elucidate these early regulators of inflammation, we performed a genome-wide CRISPR knockout screen in macrophages to identify genes that influence the induction of TNF and iNOS upon infection with Mtb. The resulting dataset is a valuable resource that includes genes representing a wide range of unexpected regulatory mechanisms that control cytokine responses to Mtb and also cell-intrinsic resistance to infection by the bacterial pathogen *Listeria monocytogenes.* We show that type I IFN signaling enhances TNF production early after infection, and IRF2 acts to inhibit induction of the antibacterial state of macrophages. Our data support a model in which early production of type I IFN in response to bacterial infection serves to increase innate antibacterial resistance during the earliest stages of infection.

## INTRODUCTION

Infection with *Mycobacterium tuberculosis* (Mtb), the causative agent of tuberculosis, is a major source of morbidity and mortality, and new therapies are needed for global tuberculosis (TB) control (Global Tuberculosis Report 2024, 2024). One approach is the use of host-directed therapies that target host processes to support eradication of Mtb. To develop these therapies, it is vital that we understand the host mechanisms that impact Mtb infection.

Mtb infection of humans results in a wide range of clinical outcomes - some people can clear early infection, some develop latent Mtb that can remain asymptomatic for decades before reactivating, while others develop symptomatic or ‘active’ TB early after infection. The mechanistic basis for this heterogeneity remains poorly understood (Simmons et al., 2018). However, it is now clear that an imbalanced host inflammatory response can drive Mtb pathogenesis. While mouse and human studies have identified several pro-inflammatory cytokines required for resisting Mtb, excessive inflammation during infection can augment Mtb growth (Flynn et al., 1995; Harris and Keane, 2010; Mayer-Barber et al., 2010; Barber et al., 2011; Mishra et al., 2017). For instance, accumulation of surplus neutrophils in the lung can result in excessive inflammation and susceptibility to infection, and depletion of neutrophils can reduce Mtb growth in many susceptible knockout mice (Nair et al., 2018; Dorhoi et al., 2010). A clearer understanding of the regulatory mechanisms involved in inflammatory responses early during infection with Mtb could improve our understanding of the varied clinical outcomes seen in human patients.

Macrophages are the first cells infected by Mtb and a major site of bacterial replication, and thus inflammatory signaling in these cells can influence the inflammatory environment in the tissue and ensuing Mtb infection. Engagement of inflammatory pathways in macrophages, including those initiated during the innate phase of bacterial infection, are vital for Mtb resistance. For example, mice lacking the receptor for the pro-inflammatory cytokine tumor necrosis factor (TNF) in macrophages, monocytes, and neutrophils demonstrate significant increases in Mtb growth *in vivo* (Segueni et al., 2016). After the onset of adaptive immunity, IFNγ activation of macrophages induces a variety of mechanisms to control Mtb, including induction of inducible expression of nitric oxide synthase (iNOS) and its product nitric oxide (NO), which can limit Mtb growth in macrophages (Chan et al., 1992). However, as inflammatory signaling pathways are subject to complex regulatory mechanisms, many of these pathways remain poorly understood. And a comprehensive understanding of the inflammatory signaling pathways that impact Mtb infection of macrophages is lacking.

Type I interferon (IFN) production and subsequent signaling through the interferon alpha/beta receptor (IFNAR) is principally associated with anti-viral immune responses and often is antagonistic to inflammatory and anti-bacterial responses. Type I IFNs dampen proinflammatory cytokine production and worsen tuberculosis disease in mice and humans (Cantaert et al., 2010; Stanley et al., 2007; Ji et al., 2021; Zhang et al., 2018). However, under some conditions IFNAR signaling is required for optimal TNF production in response to Mtb infection, and IFNAR signaling has been demonstrated to play a role in NO production upon Mtb infection (McNab et al., 2014; Moreira-Teixeira et al., 2016). More information is needed to clarify the conditions under which type I IFN responses are anti-bacterial and when they can promote bacterial growth.

Interferon regulatory factor 2 (IRF2) is a transcription factor first described as a negative regulator of the induction of type I IFN (Matsuyama et al., 1993). IRF2 has previously been implicated in the inhibition of inflammatory responses, although this regulation has been described to be independent from its regulation of type I IFN (Qin et al., 2024; Cui et al., 2018). In addition, IRF2 also directly induces expression of several innate immune genes (Kayagaki et al., 2019; Sun et al., 2017).

Here, we report the results of a genome-wide clustered regularly interspaced short palindromic repeats (CRISPR) knockout screen in Mtb-infected macrophages that identified genes involved in regulating early innate immune responses during the initial interaction of macrophages and Mtb. By examining responses generated prior to the induction of adaptive immunity, we sought to identify genes that first establish the inflammatory balance in the tissue and thereby influence later infection outcomes. Our screen identified many new factors and pathways that control early inflammatory signaling, which together serve as a fundamental resource to seed further investigation. Additionally, we show that, contrary to its antagonist role later in infection, IFNAR signaling augments pro-inflammatory and anti-bacterial responses at the early stages of infection. The transcription factor IRF2 negatively regulates many of these anti-bacterial responses, suggesting that expression levels of IRF2 may regulate the balance between anti-inflammatory versus proinflammatory and anti-bacterial responses downstream of IFNAR signaling.

## RESULTS

### Genome-wide screen for TNF and iNOS induction during Mtb infection of macrophages

We sought to explore the early innate host macrophage inflammatory response to Mtb infection by identifying the genes required for production and regulation of two key mediators induced early during infection, TNF and iNOS. Production of the proinflammatory cytokine TNF is critical for host control of Mtb infection, and NO generated by iNOS is directly antimicrobial and regulates the host inflammatory response (Flynn et al., 1995; Keane et al., 2001; Chan et al., 1992; Braverman and Stanley, 2017). To identify novel regulators of these inflammatory mediators we developed a fluorescence-activated cell sorting (FACS) based CRISPR screen examining macrophage production of TNF and iNOS upon infection with Mtb. To this end, we generated a genome-wide knockout library by transducing Cas9-expressing mouse conditionally immortalized macrophage (CIM) progenitors with the pooled sgRNA Brie library, which targets ∼20,000 genes (Roberts et al., 2019; Doench et al., 2016). The differentiated knockout library was infected with Mtb, and 24 hours post-infection the macrophages were fixed, and levels of TNF and iNOS were determined by intracellular cytokine staining. These cells were sorted into two populations, TNF^+^iNOS^-^ and TNF^+^iNOS^+^ (Fig 1A). As we sought to examine the early innate response to Mtb, the cells were not stimulated with IFNγ, therefore iNOS expression was induced in only a small proportion of macrophages. Guides from the sorted populations were amplified and sequenced and compared to those of the unsorted CIM population. Using a significance cutoff of robust rank aggregation (RRA) score < 0.001, we identified 295 genes whose knockout altered sorting into the TNF^+^iNOS^-^ population (Fig 1B), 175 were depleted and 120 enriched. We identified 184 genes whose knockout altered sorting into the TNF^+^iNOS^+^ population (Fig 1C), 81 were depleted and 103 were enriched. We identified known regulators including toll-like receptor (TLR) and NOD2 signaling pathway components, and *Tnf* itself, indicating that our approach was successful (Drennan et al., 2004; Gandotra et al., 2007).

**Figure 1:**
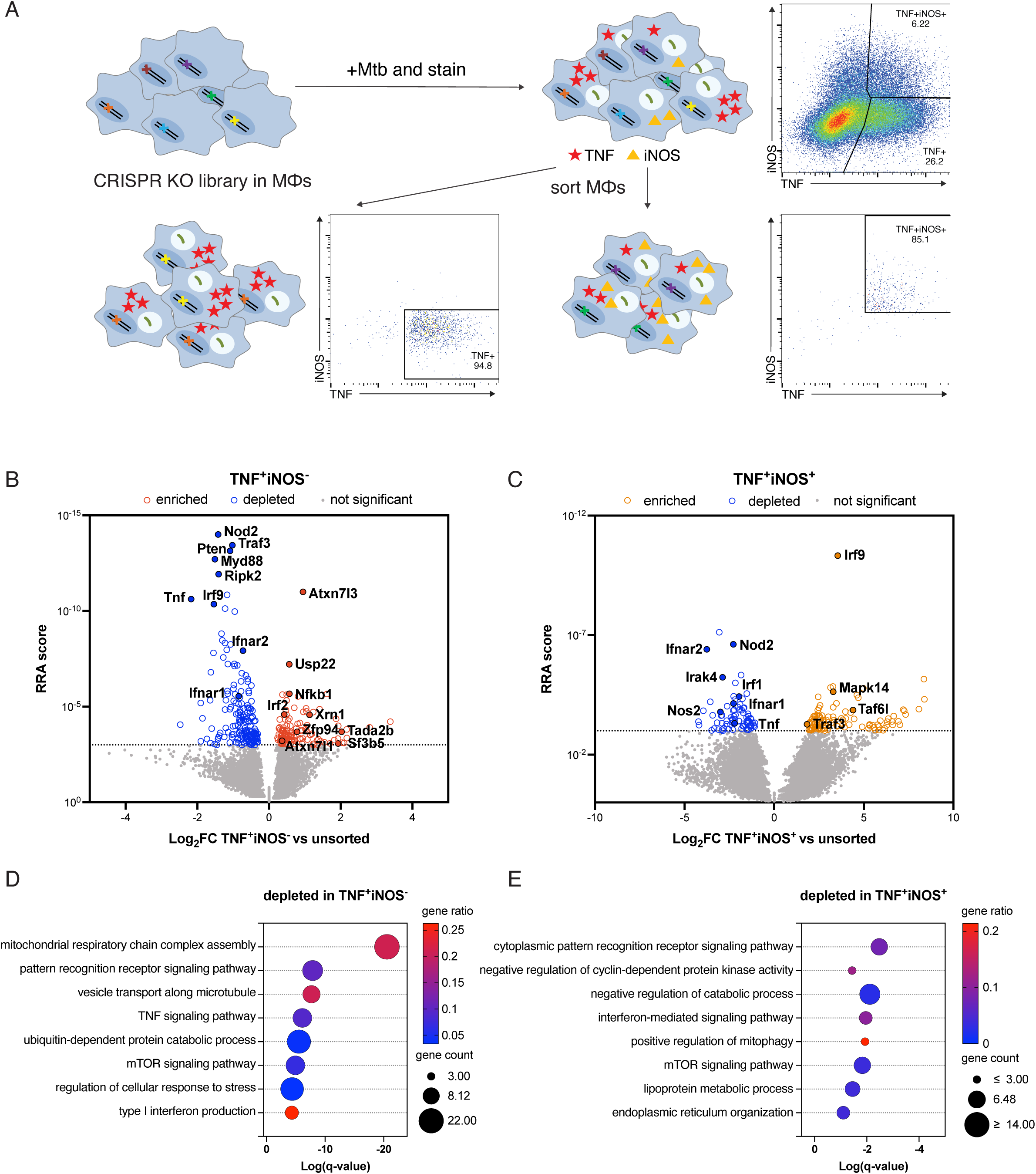
Genome-wide CRISPR knockout screen identifies regulators of TNF and iNOS in response to *Mycobacterium tuberculosis* infection of macrophages. (A) Schematic of screen. A genome-wide knockout library in CIMs was sorted based on TNF and iNOS expression during infection with Mtb. TNF and iNOS protein levels were analyzed by flow cytometry 24 hrs post infection and TNF^+^iNOS^-^ and TNF^+^iNOS^+^ CIMs were sorted as shown. (B and C) Results of genes depleted or enriched in (B) TNF^+^iNOS^-^ or (C) TNF^+^iNOS^+^ sorted macrophages compared to unsorted macrophages. (D and E) Gene set enrichment analysis of genes depleted in TNF^+^iNOS^-^ (D) or TNF^+^iNOS^+^ (E) macrophage populations. Data are the combined results of two independent experiments.

Pathway analysis of the genes in these two sorted populations revealed several notable gene sets that were significantly enriched in the data (Fig 1D and 1E). Genes involved in TNF signaling and pattern recognition receptor signaling were depleted in the TNF^+^iNOS^-^ population, indicating that our screening methodology was able to identify known TNF regulators. By far the most significantly enriched gene set was the mitochondrial respiratory chain complex assembly pathway, of which we identified 19 genes encoding components of respiratory chain complex I that were highly depleted in the TNF^+^iNOS^-^ population. Mitochondria-derived reactive oxygen species generated by perturbations in complex I have been linked to regulation of macrophage inflammation and IL-1β production (Carneiro et al., 2018; Mills et al., 2016; Casey et al., 2025). Our data provides compelling evidence that complex I is required for full production of TNF upon infection with Mtb, however the mechanisms involved remain unclear.

We also identified components of the transcriptional coactivator Spt-Ada-Gcn5 acetyltransferase (SAGA) complex, *Atxn7l1, Atxn7l3, Tada2b, Usp22, Sf3b5*, and *Taf6l*, which were enriched in the TNF^+^iNOS^-^ or TNF^+^iNOS^+^ populations. This complex promotes Foxp3 protein expression in regulatory T cells, but to our knowledge an inhibitory role in TNF production has not previously been demonstrated (Loo et al., 2020).

Due to previously described antagonism between type I IFN and inflammatory responses (Cantaert et al., 2010; Feng et al., 2024) we were surprised to find that genes required for IFNAR signaling, including the receptor genes *Ifnar1* and *Ifnar2*, as well as transcription factor *Irf9*, were depleted in the TNF^+^iNOS^-^ population, suggesting that type I IFN promotes TNF production under these conditions (Figs 1B-1E).

Consistent with this notion, *Irf2*, a negative regulator of type I IFN production, was enriched in the TNF^+^iNOS^-^ population (Matsuyama et al., 1993)(Fig 1B). We also identified IFNAR pathway components that were depleted in in the TNF^+^iNOS^+^ population, which indicates a role in iNOS induction as well.

### Secondary arrayed screen identifies novel regulators of anti-bacterial responses

To further investigate the genes identified by the CRISPR screen we performed an arrayed secondary screen in which ten candidate genes were individually knocked out in CIMs using two independent guides. We infected macrophages lacking the candidate gene with fluorescent Mtb, and analyzed Mtb growth as well as TNF and NO levels (Fig 2 A-C). As a control, knockout of the genes *Myd88* and *Nod2*, which are required for complete TNF production during Mtb infection, resulted in decreased TNF production. Treatment of macrophages with IFNγ was also included as a positive control and showed augmented production of TNF and NO, as well as restriction of Mtb growth. Although no candidate genes dramatically altered growth of Mtb, we were able to confirm that many of these genes regulate TNF and/or NO production. TNF production was reduced in macrophages lacking *Ifnar1*, *Irf9*, and *Traf3* (Fig 2B) demonstrating that IFNAR signaling is needed for optimal production of TNF upon Mtb infection. However, although *Traf3* has been shown to promote type I IFN production, it’s unclear if the observed decrease in TNF production in macrophages lacking *Traf3* is due to type I IFN responses as TRAF3 plays a role in a variety of signaling pathways (Oganesyan et al., 2006; Lin et al., 2023). Interestingly, we also identified two candidate genes, *Atxn7l3* and *Irf2*, whose knockout enhanced TNF production in response to Mtb infection.

**Figure 2:**
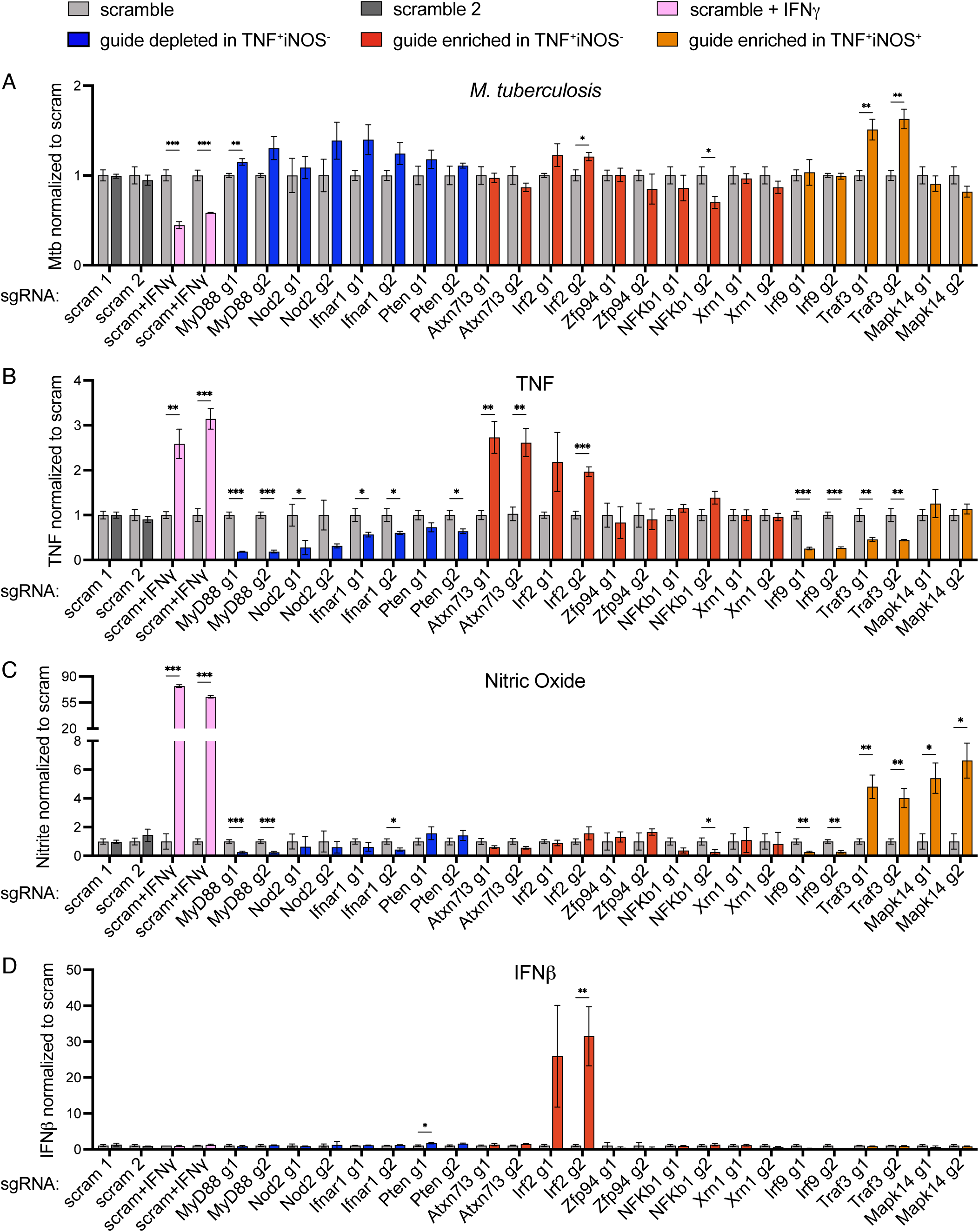
Arrayed secondary screen identifies regulators of cytokine and nitric oxide induction during *Mycobacterium tuberculosis* infection of macrophages. CIMs lacking candidate genes were generated using two independent sgRNA for each candidate, and infected with fluorescent Mtb. (A) Four days post infection Mtb growth in macrophages was analyzed by quantifying fluorescent area in KO macrophages normalized to fluorescent area in control (scramble gRNA) macrophages. (B-D) Two days post infection levels of (B) TNF, (C) nitric oxide, and (D) IFNβ were measured and normalized to levels generated by control macrophages. Graphs show compiled data from four experiments, each examining different guides and the corresponding controls. Data for each guide show the mean +/- SD from one experiment with four replicates. A second independent experiment confirmed the results for *Ifnar1, Atxn7l3, Irf2,* and *Traf3*. Data were statistically analyzed using t-test with Holm-Sidak’s multiple comparisons test. P < 0.05, **P < 0.01, ***P < 0.001, **** P < 0.0001.

Finally, macrophages lacking two of the genes identified in the TNF^+^iNOS^+^ sorted population, *Traf3* and *Mapk14*, exhibited enhanced production of NO upon Mtb infection. Overall, the secondary screen revealed several novel regulators of TNF and NO production during Mtb infection.

We analyzed the production of several other important cytokines induced by Mtb. We observed a dramatic increase in IFNβ production in macrophages lacking *Irf2* (Fig 2D). IRF2 has previously been described to inhibit type I IFN production, and these data demonstrate this activity is vital for regulating type I IFN production during Mtb infection (Matsuyama et al., 1993). We also measured differences in IL-6 and IL-1α production in macrophages lacking several of the candidate genes (Fig S1 A and B). Most notably, macrophages lacking *Pten* generated significantly more IL-1α in response to Mtb infection. Further investigation is required to determine whether these cells generate higher levels of IL-1α protein or if macrophages lacking *Pten* are more likely to die during Mtb infection, leading to enhanced IL-1α release (Cavalli et al., 2021).

Although none of the candidate genes impacted Mtb growth in knockout macrophages, Mtb is an exceptionally resilient pathogen that is resistant to nearly all anti-microbial macrophage functions induced during the early innate immune response (Chandra et al., 2022). Thus, to test the potential anti-bacterial mechanisms regulated by these genes, we infected macrophages with the opportunistic intracellular bacterial pathogen *Listeria monocytogenes* (Lm), which also replicates in macrophages. After phagocytosis, Lm quickly escapes from the phagosome into the host cytosol where it replicates rapidly. CIMs lacking candidate genes were infected with fluorescent Lm and five hours post infection Lm growth and cytokine production were analyzed (Fig 3 A-C). As shown in Figure 3A, the growth of Lm was significantly restricted in macrophages lacking *Irf2*. This growth restriction is further explored in Figures 6 and 7 (see below).

**Figure 3:**
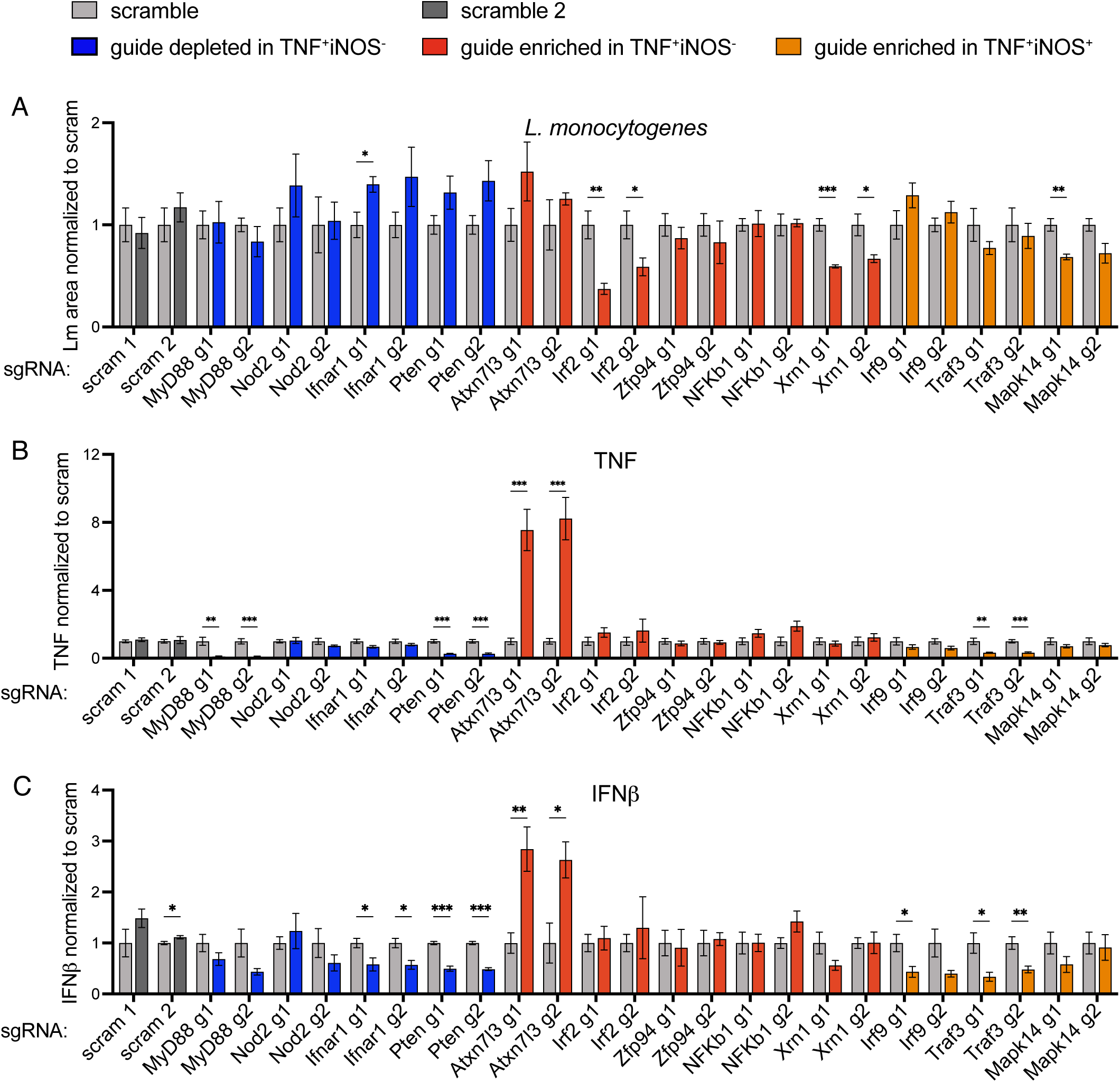
Arrayed secondary screen identifies regulators of bacterial growth and cytokine responses during *Listeria monocytogenes* infection of macrophages. CIMs lacking candidate genes were generated using two independent sgRNA for each candidate, and infected with fluorescent Lm. (A) Five hours post infection Lm growth in macrophages was analyzed by quantifying fluorescent area in KO macrophages normalized to fluorescent area in control (scramble gRNA) macrophages. (B-C) Cell supernatants were measured for levels of (B) TNF and (C) IFNβ and normalized to levels generated by control macrophages. Graphs show compiled data from four experiments, each examining different guides and the corresponding controls. Data for each guide show the mean +/- SD from one experiment with four replicates. A second independent experiment confirmed the results for *Ifnar1, Pten, Atxn7l3, Irf2, Traf3,* and *Mapk14*. Data were statistically analyzed using t-test with Holm-Sidak’s multiple comparisons test. *P < 0.05, **P < 0.01, ***P < 0.001, **** P < 0.0001.

Although we did not measure restriction of Mtb growth in macrophages lacking *Irf2*, we did find enhanced production of TNF, indicating that IRF2 inhibits induction of anti- microbial responses during both infections.

Many of the observed cytokine response phenotypes generated by macrophages lacking our candidate genes were similar in Mtb and Lm infection. For instance, during infection with Mtb and Lm macrophages lacking *Atxn7l3* generated significantly more TNF, while macrophages lacking *Traf3* generated less TNF (Fig 2B and 3B).

Additionally, IL-6 production was significantly reduced in macrophages lacking *Ifnar1* and *Irf9*, components of IFNAR signaling, during both Mtb and Lm infections (Fig S1A and S1C).

There were also some notable differences in the cytokine responses generated upon Mtb and Lm infection. For instance, while both *Myd88* and *Nod2* were vital for TNF responses to Mtb infection, *Nod2* deficiency did not impact TNF responses to Lm. This result is consistent with previous studies demonstrating that although *Nod2* KO impacts Lm infection in the intestine, it does not impact TNF produced by BMMs upon Lm infection (Kobayashi et al., 2005). This seems to be due to a bacterial evasion mechanism in which Lm modifies its peptidoglycan to reduce susceptibility to lysozyme and evade recognition by NOD2 (Boneca et al., 2007). In contrast, Mtb modifies its peptidoglycan in a manner that also increases resistance to lysozyme but enhances recognition by NOD2 receptors (Raymond et al., 2005; Coulombe et al., 2009). In addition, during infection with Lm macrophages, lacking *Irf2* did not generate the robust TNF and IFNβ production seen during Mtb infection, perhaps due to reduced Lm growth, and thus reduced activation of pattern recognition receptors (PRRs).

### *Traf3*-deficient macrophages generate reduced TNF responses and modestly restrict Lm growth

We further investigated the inflammatory responses and anti-bacterial activity of macrophages lacking *Traf3*. We measured reduced TNF levels in *Traf3*-deficient macrophages during both Mtb and Lm infections but enhanced NO production during Mtb infection (Fig 2B, 2C, and 3B). It has been reported that while *Traf3*-deficient macrophages generate less type I IFN, they produce similar levels of TNF after LPS stimulation (Lalani et al., 2015). To test whether the reduction in TNF we measured was specific to bacterial infection, we stimulated *Traf3*-deficient CIMs and bone marrow derived macrophages (BMMs) with the TLR2 ligand Pam3CSK4, the TLR9 ligand CpG, or the NOD2 ligand MDP, with and without the addition of IFNβ. In both CIMs and BMMs we measured dramatically reduced production of TNF in *Traf3*-deficient macrophages after stimulation with TLR2 or TLR9 ligands (Fig S2A and C). When stimulated with MDP + IFNβ we observed enhanced production of NO in *Traf3*-deficient macrophages (Fig S2B and D). These results correlated with the responses we observed during Mtb infection, though the mechanism of TRAF3 regulation of TNF and NO production remains unclear. Although TRAF3 is required for maximal induction of type I IFN after stimulation of several PRRs, differences in type I IFN production are likely not responsible for the observed TNF and NO induction phenotypes, as addition of IFNβ does not meaningfully increase TNF induction in the *Traf3*-deficient cells (Fig S2A and C). We also investigated whether TRAF3 regulated anti-bacterial mechanisms, as we noted a small trend towards reduced Lm growth in *Traf3*-deficient macrophages in our secondary screen. By both microscopic and CFU analysis we observed a small but statistically significant reduction in Lm growth in *Traf3*-deficient BMMs (Fig S2E and F). TNF production in the *Traf3*-deficient BMMs was also dramatically reduced during Lm infection compared to controls (Fig S2E).

### IRF2 inhibits production of TNF

Our finding that IFNAR signaling is required for optimal induction of TNF and iNOS during Mtb infection led us to hypothesize that innate immune cell-generated type I IFN can induce pro-inflammatory and anti-bacterial mechanisms to restrain bacterial growth before the adaptive response is activated. To test this, we examined whether IFNAR signaling can augment TNF and NO production in response to TLR ligand stimulation. Stimulation of BMMs with TLR9 or TLR2 ligands resulted in reduced TNF production in IFNAR-deficient BMMs, and addition of IFNβ to WT BMMs significantly increased production of both TNF and NO compared to ligand alone (Fig 4A and 4B).

**Figure 4:**
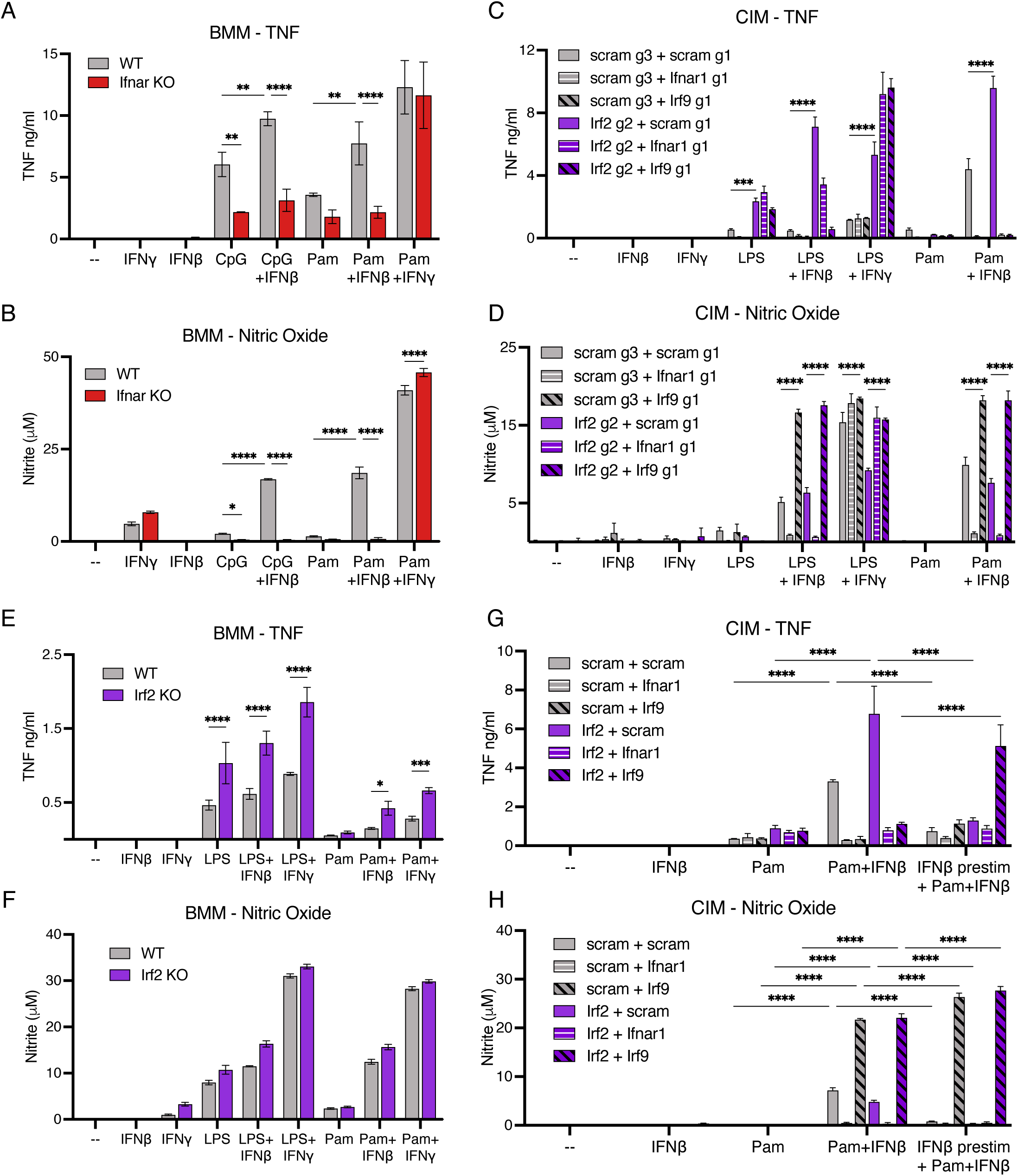
IRF2 inhibits TNF production in response to PRR stimulation. (A) TNF and (B) nitric oxide production by WT and IFNAR KO BMMs stimulated with CpG or Pam3CSK4 +/- IFNβ or IFNγ. Data are representative of at least two independent experiments. (C) TNF and (D) Nitric oxide production by double knockout CIMs (transduced with guides for *Irf2* or scramble and *Irf9, Ifnar1,* or scramble) stimulated with LPS or Pam3CSK4 +/- IFNβ or IFNγ. Data are representative of at least three independent experiments. (E) TNF and (F) nitric oxide production by WT and *Irf2* KO BMMs stimulated with LPS or Pam3CSK4 +/- IFNβ or IFNγ. Data are representative of at least three independent experiments. (G) TNF and (H) nitric oxide production by double knockout CIMs prestimulated +/- IFNβ for 24 hours before stimulation with Pam3CSK4 +/- IFNβ. Data are representative of at least three independent experiments. All data are presented as mean +/- SD and were statistically analyzed using two-way ANOVA with Sidak’s multiple comparisons test. *P < 0.05, **P < 0.01, ***P < 0.001, **** P < 0.0001.

This result correlates with the results from our Mtb infections (Fig 2), as well as previous studies demonstrating that IFNAR signaling can boost production of NO and TNF in response to Mtb infection (Moreira-Teixeira et al., 2016; McNab et al., 2014). Because IRF1 expression is induced by IFNAR signaling and is required for iNOS induction downstream of IFNγR signaling (Kamijo et al., 1994), we tested whether this factor is responsible for TNF and NO production in response to type I IFN. CIMs lacking *Irf1* induced significantly less NO upon stimulation with Pam3CSK4 or MDP plus IFNβ (Fig S3B). However, the increase in TNF measured in response to IFNβ was not dependent on IRF1 (Fig S3A).

To determine whether IFNAR-induced inflammatory responses are regulated by IRF2, we generated CIMs deficient for both *Irf2* and either *Irf9* or *Ifnar1*. We used two cas9 guides for each gene to generate two independent double knockout (DKO) macrophage lines for each combination. When stimulated with the TLR4 ligand LPS, *Irf2*-deficient macrophages generated significantly heightened TNF levels (Fig 4C and Fig S3C). This augmented TNF production was not dependent on IFNAR signaling as *Irf2* x *Ifnar1* DKO and *Irf2* x *Irf9* DKO cells also generated more TNF in response to LPS. However, co-administration of IFNβ with either LPS or Pam3CSK4 further increased TNF production in *Irf2* KO macrophages in an IFNAR-dependent manner. We did not detect dramatic differences in NO production between controls and *Irf2*-deficient macrophages, though macrophages lacking *Irf9* generated significantly more NO (Fig 4D and S3D). This result correlates with our CRISPR screen in which *Irf9* was enriched in the TNF^+^iNOS^+^ population, while *Ifnar1* and *Ifnar2* were depleted. The canonical IFNAR signaling pathway induces formation of a transcription factor complex containing IRF9, signal transducer and activator of transcription 1 (STAT1), and STAT2, however other complexes, including homodimers of STAT1, can be induced downstream of IFNAR engagement that do not contain IRF9. We suspect that the increase in NO seen in *Irf9*-deficient macrophages is due to enhanced formation of STAT1 homodimers, which has previously been demonstrated to occur in *Irf9*-deficient cells (Gothe et al., 2022). To test whether these effects are manifest in primary macrophages, we stimulated BMMs derived from *Irf2* KO mice (Kayagaki et al., 2019). As observed in CIMs, we measured enhanced production of TNF, but not NO, in *Irf2* KO BMMs after stimulation with LPS or Pam3CSK4 plus IFNβ (Fig 4E and 4F). A recent publication noted IRF2 mediated inhibition of IL-6 and iNOS responses after stimulation with LPS + IFNγ, and suggested that this inhibition is dependent on IRF2 mediated expression of IRG1 (Qin et al., 2024). However, TNF and NO levels were largely unaffected in CIMs lacking both *Irf2* and *Irg1* compared to single *Irf2-*deficient cells, indicating that IRG1 is not responsible for the phenotype (Fig S3E and S3F).

In the experiments described above we added TLR ligands and IFNβ simultaneously since type I IFN production during a bacterial infection would not precede PRR stimulation. We wondered whether IFNβ exposure prior to TLR stimulation would inhibit IFNAR induced TNF and NO production and if IRF2-mediated regulation of TNF would be impacted. Indeed, when pretreated with IFNβ, control and *Irf2*-deficient macrophages produce less TNF and NO compared to cells that had received both treatments simultaneously (Fig 4G and 4H). Interestingly, although IFNβ- induced augmentation of TNF after simultaneous co-stimulation required IRF9, IFNβ pretreatment induced augmented TNF production only in *Irf2* x *Irf9* DKO macrophages. These data suggest that with IFNβ prestimulation IRF9-independent noncanonical IFNAR signaling can induce TNF production. TNF induction by this noncanonical signaling is also inhibited by IRF2. Overall, the timing of IFNβ addition dramatically alters TNF and NO production, and is one possible source of the varied results reported in the literature regarding the impact of IFNAR signaling on proinflammatory responses (Ji et al., 2023).

### IRF2 inhibits proliferation during macrophage differentiation

Enhanced monocyte proliferation can enable rapid cell recruitment to combat infection. For instance, emergency myelopoiesis in the bone marrow can be induced by a variety of cytokines and PRR signals to increase monocyte and neutrophil numbers (Orozco et al., 2021). Recruited monocytes have also been reported to undergo local proliferation prior to differentiation into tissue resident macrophages (Vanneste et al., 2023). We observed a dramatic increase in cell number after differentiation of *Irf2* x *Irf9* and *Irf2* x *Ifnar1* DKO CIMs (Fig 5A). This increased proliferation did not occur in macrophages lacking either *Irf2* or components of the IFNAR signaling pathway alone.

**Figure 5:**
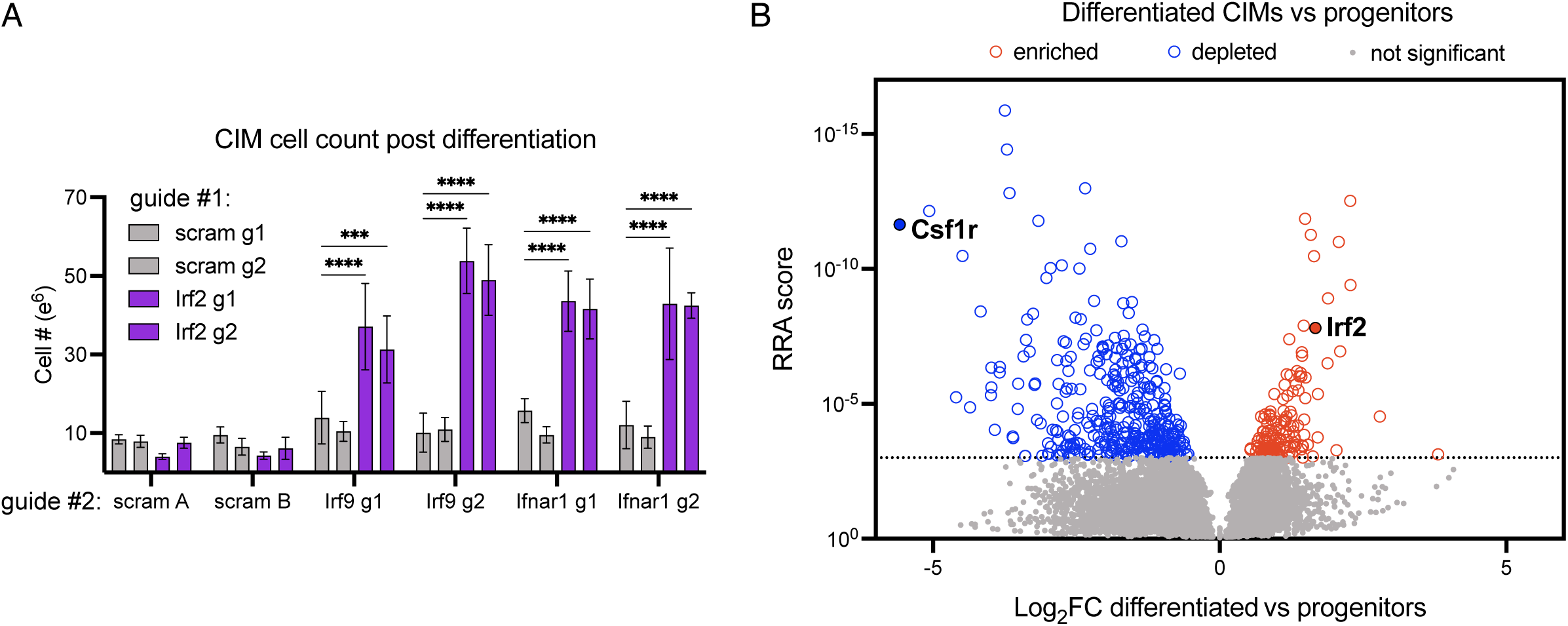
IRF2 inhibits proliferation during macrophage differentiation. (A) 2x10^6^ double knockout CIM progenitors (transduced with guides for *Irf2* or scramble and *Irf9, Ifnar1*, or scramble) were differentiated for seven days and cell numbers were quantified. Data are the combined results of at least three independent experiments. Data are presented as mean +/- SD and were statistically analyzed using two-way ANOVA with Sidak’s multiple comparisons test. *P < 0.05, **P < 0.01, ***P < 0.001, **** P < 0.0001. (B) Genome wide CRISPR knockout screen examining macrophage differentiation and proliferation. The CIM progenitor knockout library was differentiated into macrophages, and undifferentiated cells at day 0 were compared to cells differentiated for seven days. Shown are results of genes depleted or enriched in differentiated CIMs compared to progenitors. Data are the combined results of three independent experiments.

As IFNAR signaling is known to decrease cell proliferation, we wondered whether slightly enhanced type I IFN levels produced by *Irf2*-deficient cells during differentiation may limit cell proliferation and obscure IRF2-mediated regulation of proliferation in cells with intact IFNAR signaling. To investigate the impact of *Irf2* deficiency without the added complication of heightened type I IFN signaling, we performed a differentiation screen using our genome-wide CIM knockout library. As the library includes mostly cells that express IRF2, any increase in type I IFN produced by extremely rare *Irf2*-deficient cells should be inconsequential in the population. To identify genes involved in macrophage differentiation, we compared sgRNA guides in the progenitor library to those in differentiated CIMs. In the presence of β-estradiol, self-renewing CIM progenitors remain undifferentiated and non-adherent, but removal of estradiol and addition of the growth factor macrophage colony-stimulating factor (MCSF) promotes differentiation into macrophages that adhere tightly to polystyrene culture flasks (Wang et al., 2006; Roberts et al., 2019). Seven days after induction of differentiation, macrophages were selected based on adherence and guide libraries were sequenced. To identify genes specifically required for macrophage differentiation, we compared these guide sequences to sequences generated from the starting stock of undifferentiated CIMs, as well as from cells maintained as progenitors and passaged in parallel to the differentiated macrophages. In both comparisons, we found that *Csf1r*, a gene vital macrophage differentiation, was depleted, demonstrating that the screen was able to identify known factors (MacDonald et al., 2010). Importantly, *Irf2* guides were enriched in differentiated macrophages suggesting that cells lacking IRF2 proliferated more during macrophage differentiation (Fig 5A and S4A). Collectively, these data indicate that IRF2 also regulates cell proliferation during macrophage differentiation.

Since the differentiation of CIMs can be precisely controlled by removal of estradiol and addition of MCSF, our dataset is a well-controlled survey of the genes specifically involved in differentiation and proliferation of macrophages and eliminates genes that are required for proliferation of progenitor cells. To affirm this approach, we individually knocked out six candidate genes using two independent guides each: three that were enriched in the differentiated macrophages and three that were depleted.

Macrophages deficient in *Morc3*, *Cul5*, and *Trps1*, that were depleted in the differentiation screen, demonstrated drastically lower cell counts. After differentiation we were unable to recover enough cells deficient in *Morc3* for analysis, but cells deficient for *Cul5* and *Trps1* exhibited reduced expression of the macrophage marker F4/80, suggesting they did not properly differentiate (Fig S4B and C). Macrophages lacking *Adrbk1* and *Mapk7,* two of the three genes that were enriched in the differentiated population, demonstrated enhanced cell counts post differentiation. These data support a previous study that demonstrated *Mapk7* activity inhibits macrophage-like phenotypes in human acute myeloid leukemia cells (Wang et al., 2015). To our knowledge, the other genes identified in the secondary screen have not previously been shown to impact macrophage differentiation. Thus, this dataset provides a valuable resource for understanding additional mechanisms of macrophage differentiation and proliferation.

### Restriction of Lm growth in *Irf2* KO macrophages is dependent on IFNAR signaling

To elucidate the anti-bacterial responses regulated by IRF2, we further explored the mechanisms mediating the restriction of Lm growth in *Irf2*-deficient macrophages. First, we confirmed the bacterial growth restriction by enumeration of wild type Lm CFU in both CIMs and BMMs, and measured between two and four fold restriction of Lm growth in macrophages lacking *Irf2* (Fig 6A and 6B). Lm growth in *Irf2* KO macrophages was restored to normal levels in *Irf2* x *Ifnar1* DKO and *Irf2* x *Irf9* DKO cells, demonstrating that restriction was due to induction of anti-microbial mechanisms downstream of IFNAR signaling (Fig 6C and 6D). As several different transcription factor complexes have been shown to function downstream of IFNAR engagement, we sought to determine if the canonical complex containing IRF9, STAT1 and STAT2 is required. We found that all components of this complex are required for Lm growth restriction in *Irf2*-deficient macrophages (Fig 6E). Although we did not measure enhanced production of IFNβ in *Irf2*-deficient macrophages during Lm infection (Fig 3C), we sought to remove the possibility that heightened type I IFN levels are the source of Lm restriction in *Irf2*-deficient macrophages. Therefore, we ensured that all macrophage genotypes received significant type I IFN by pretreating macrophages with IFNβ prior to infection with Lm. Post IFNβ pretreatment, we continued to measure reduced Lm growth in *Irf2*-deficient macrophages relative to controls (Fig 6F). Interestingly, we also observed Lm growth restriction in *Irf9*-deficient macrophages independent of Irf2 expression, suggesting that induction of a non-canonical pathway downstream of IFNAR engagement can also induce anti-bacterial mediators that restrict Lm growth. Overall, these data demonstrate that IRF2 inhibits an antimicrobial mechanism that can be induced by canonical IFNAR signaling.

**Figure 6:**
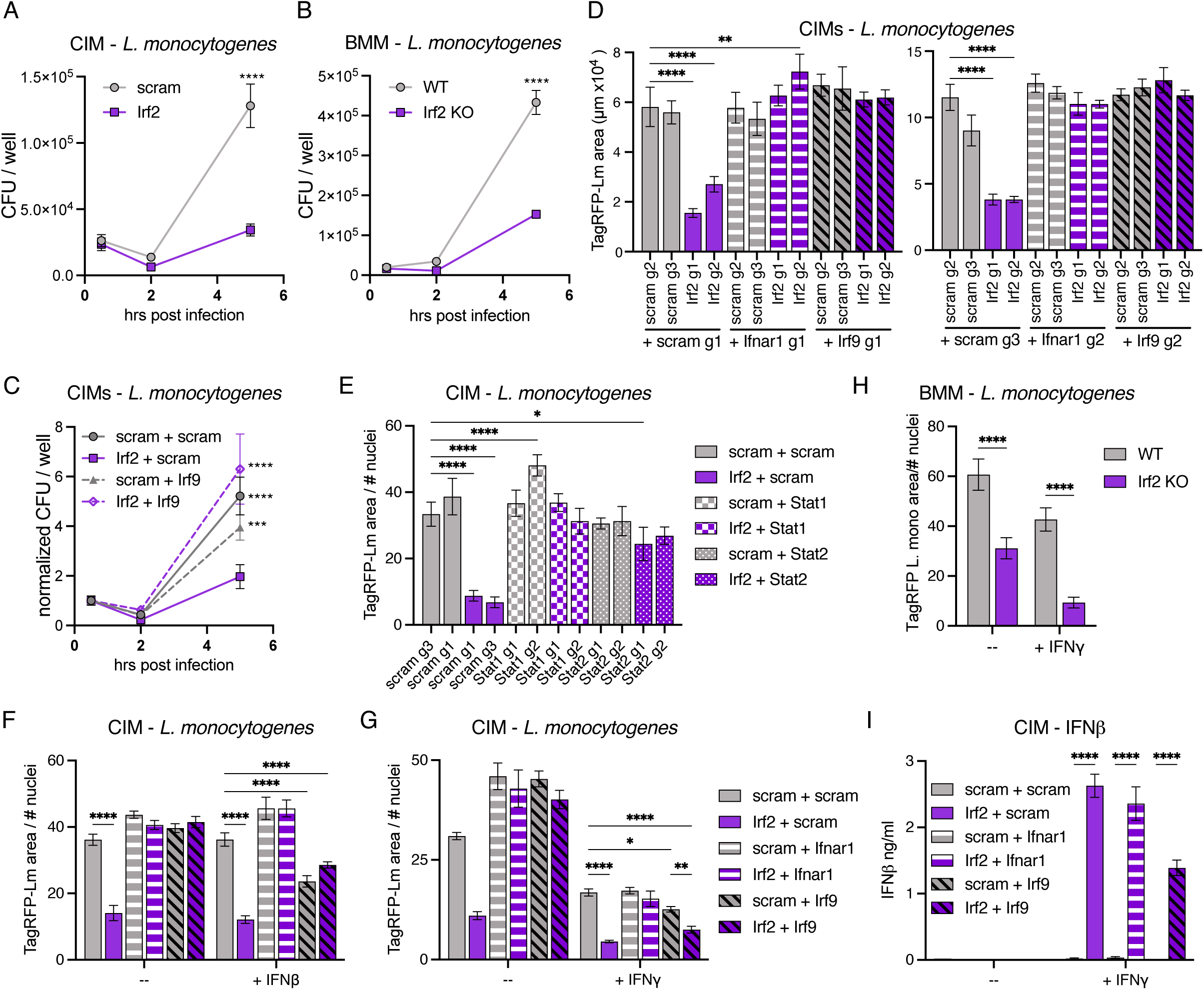
*Listeria monocytogenes* growth is restricted in *Irf2* deficient macrophages downstream of IFNAR signaling. (A-C) WT Lm intracellular CFU were enumerated at indicated time points post infection in (A) CIMs, transduced with guides for *Irf2* or scramble controls, (B) control and *Irf2* KO BMMs, or (C) double knockout CIMs (transduced with guides for *Irf2* or scramble and *Irf9* or scramble). Data are representative of at least three independent experiments. (D) Double knockout CIMs (transduced with guides for *Irf2* or scramble and *Ifnar1, Irf9,* or scramble) were infected with TagRFP-Lm. Five hours post infection Lm growth in CIMs was analyzed by quantifying TagRFP area. Data are representative of at least three independent experiments. (E) Double knockout CIMs, (transduced with guides for *Irf2* or scramble and *Stat1, Stat2,* or scramble) were infected with TagRFP-Lm. Lm growth was analyzed five hours post infection. Data are representative of at least two independent experiments. (F-G) Double knockout CIMs (transduced with guides for *Irf2* or scramble and *Ifnar1, Irf9,* or scramble) were prestimulated +/- (F) IFNβ or (G) IFNγ for 18 hours before infection with TagRFP-Lm. Lm growth was analyzed five hours post infection. Data are representative of at least three independent experiments. (H) Control and *Irf2* KO BMMs were prestimulated +/- IFNγ for 18 hours before infection with TagRFP-Lm. Lm growth was analyzed five hours post infection. Data are representative of at least three independent experiments. (I) Double knockout CIMs were stimulated +/- IFNγ for 20 hours before IFNβ levels were quantified. Data are representative of at least two independent experiments. All data are presented as mean +/- SD and were statistically analyzed using one-way (D) or two-way ANOVA with Sidak’s multiple comparisons test. *P < 0.05, **P < 0.01, ***P < 0.001, **** P < 0.0001.

Because we observed heightened TNF production in *Irf2*-deficient cells stimulated with LPS and IFNγ (Fig 4C), we hypothesized Irf2 may also inhibit an antimicrobial mediator induced by IFNγR. Pretreatment of CIMs and BMMs with IFNγ prior to infection with Lm led to even greater restriction in IFNγ treated *Irf2* KO cells than control cells (Fig 6G and H). In addition, after IFNγ treatment we observed secretion of IFNβ in macrophages lacking *Irf2* but not in scramble controls (Fig 6I). In conclusion, these data suggest that downstream of IFNγR signaling IRF2 inhibits expression of type I IFN, and possibly other antimicrobial mediators.

### Restriction of Lm growth in *Irf2* KO macrophages is not dependent on IRF1, IRG1, or cell death

We next sought to identify the IRF2-regulated factors responsible for Lm restriction. Because IRF2 inhibits IRF1 mediated transcription, we hypothesized that IRF1 activation was responsible for the effects of IRF2 inactivity (Matsuyama et al., 1993; Parrington et al., 1993). However, Lm restriction in *Irf2*-deficient macrophages was not dependent on IRF1 as we measured Lm restriction in *Irf2* x *Irf1* DKO CIMs (Fig S5A). Likewise, IRG1 expression and its subsequent anti-inflammatory effects have been described to be dependent on IRF2 (Qin et al., 2024). However, *Irg1* deficiency did not alter Lm restriction in *Irf2*-deficient macrophages (Fig S5B). We also determined that neither IRF1 nor IRG1 are likely to be responsible for Lm restriction downstream of IFNγR, as Lm growth in IFNγ stimulated CIMs lacking these genes was similar to controls. Finally, as IRF2 is known to regulate pyroptosis via expression of GSDMD, we also measured macrophage death during Lm infection (Kayagaki et al., 2019). We did not observe differences in cell lysis between *Irf2*-deficient and control macrophages (Fig S5C). Therefore, we ruled out these previously described functions of IRF2 as responsible for the Lm growth restriction observed in *Irf2* KO macrophages.

### IRF2 regulates expression of antimicrobial mediators

To identify genes regulated by IRF2 and IFNAR during Lm infection, we performed RNAseq on RNA harvested from Lm infected *Irf2*, *Irf9*, and *Irf2* x *Irf9*-deficient CIMs as well as scramble controls (Fig 7A). Approximately the same number of genes were positively regulated by IRF2 as were inhibited, although without further studies we cannot determine whether this regulation is direct. IRF2 has previously been described to positively regulate *Gsdmd* and *Tlr3* expression, and we measured lower expression of these genes in macrophages lacking *Irf2* (Kayagaki et al., 2019; Sun et al., 2017).

**Figure 7:**
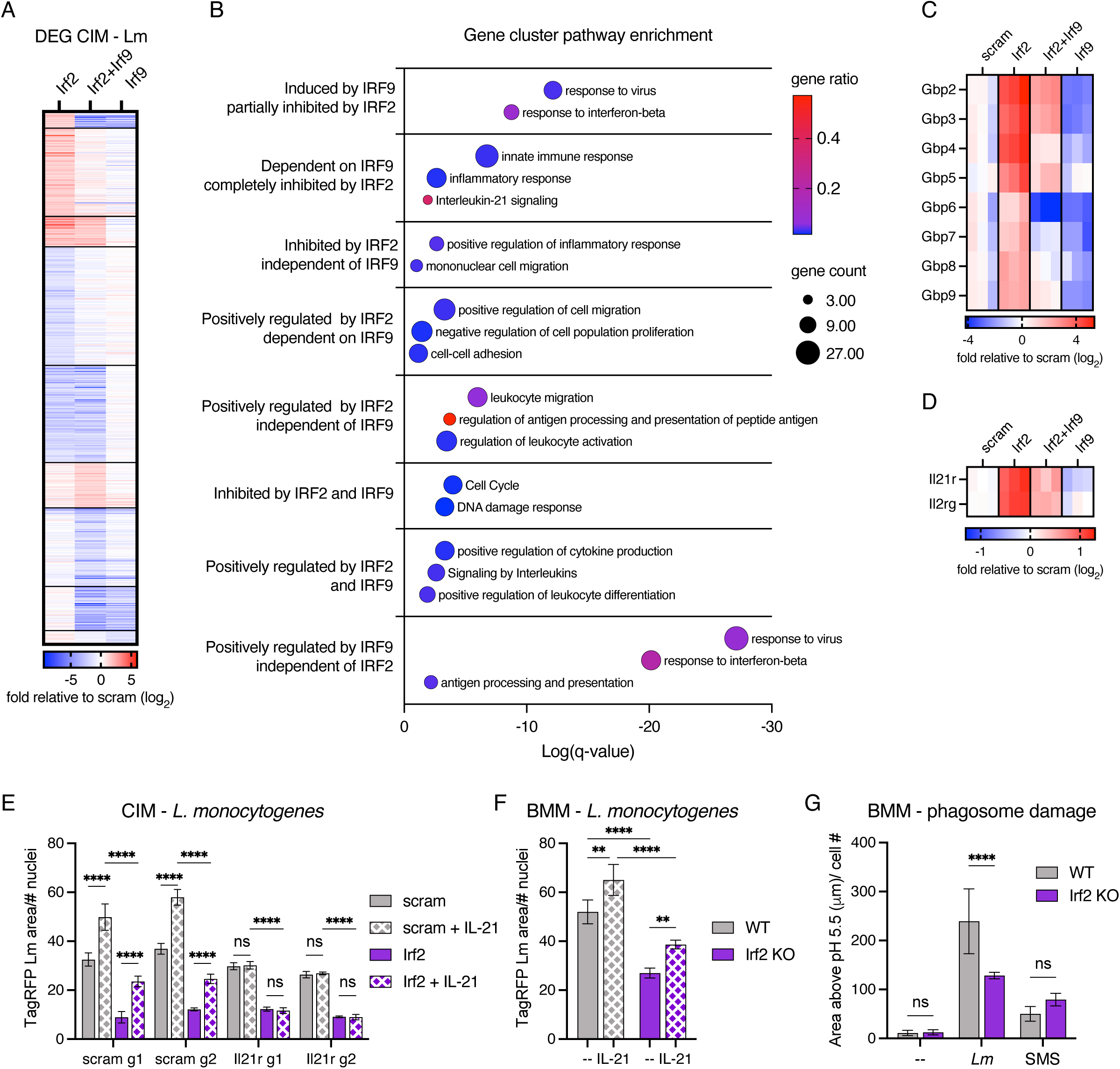
IRF2 regulates expression of antimicrobial mediators. (A-D) Double knockout CIMs (transduced with guides for *Irf2* or scramble and *Irf9* or scramble) were infected with WT Lm. Two hours post infection total RNA was collected and used for RNAseq analysis. Data are from three independent experiments. (A) Displayed are the average log2 fold change, relative to scramble, of genes with differential expression in at least one knockout macrophage line. Genes are clustered based on fold change in different knockout macrophage lines. (B) Pathway enrichment of gene clusters. (C-D) Heat map of (C) GBP family or (D) *Il21r* and *Il2rg* gene expression in knock out macrophages relative to scramble. Shown are values from three independent experiments. (E) Double knockout CIMs (transduced with guides for *Irf2* or scramble and *Il21r* or scramble) or (F) WT and *Irf2* KO BMMs were prestimulated +/- IL-21 before infection with TagRFP-Lm. Five hours post infection Lm growth was analyzed by quantifying TagRFP area. Data are representative of two (E) or three (F) independent experiments. (G) BMMs loaded with fluorescein-dextran were infected with WT Lm or administered silicon dioxide microspheres (SMS) for two hours. Fluorescence was measured after excitation at 425nm and 488nm and used to determine the area with pH above 5.5. Data are representative of two independent experiments. All data are presented as mean +/- SD and were statistically analyzed using two-way ANOVA with Sidak’s multiple comparisons test. *P < 0.05, **P < 0.01, ***P < 0.001, **** P < 0.0001.

Since Lm restriction in *Irf2* KO macrophages is dependent on *Irf9*, we compared *Irf2* KO to *Irf2* x *Irf9* DKO cells to identify the subset of IRF2-regulated genes that was also dependent on IRF9-mediated type I IFN signaling. Gene clustering based on positive or negative regulation by IRF2 and IRF9 demonstrated that both IRF2 inhibited and positively regulated genes had IRF9 dependent and independent subsets (Fig 7A). As expected, we identified a cluster of genes which demonstrate lower expression in both *Irf9* KO and *Irf2* x *Irf9* DKO cells and enhanced expression in *Irf2* KO cells, suggesting that they are induced by IRF9 and inhibited by IRF2. But, we were intrigued to see another cluster of genes whose expression was higher in *Irf2* KO cells relative to scramble, but was similar to scramble in *Irf9* KO or *Irf2* x *Irf9* DKO cells. This suggests that while IRF9 is capable of inducing expression of these genes, this IRF9-driven expression is typically prevented by IRF2.

Our RNAseq data provide new insights into how IRF2 and IRF9 interact to regulate transcription. Using gene pathway enrichment of the gene clusters, we found that many of the genes that are induced by IRF9 and partially inhibited by IRF2 were involved in responses to virus and IFNβ, which is unsurprising for IRF9-driven genes. However, genes that are dependent on IRF9 but completely inhibited by IRF2 were instead enriched for innate immune responses and inflammatory responses (Figure 7B).

This suggests that in addition to the previously described role of IRF2 in inhibiting IFNβ production, IRF2 inhibits inflammatory genes that can be induced by IRF9. Not all IRF2 inhibited inflammatory genes are dependent on IRF9; genes inhibited by IRF2 in an IRF9 independent manner were also enriched for positive regulation of inflammatory responses. Finally, we noted enrichment of cell cycle genes in the cluster of genes that are higher in only *Irf2* x *Irf9* DKO cells, this correlates with the cell proliferation phenotype we observed in *Irf2*-deficient cells that also lacked IFNAR signaling components (Fig 5A). Overall, the RNAseq analysis revealed a complex network of genes that are regulated by IRF2, many of which have pro-inflammatory functions, and a subset of which are dependent on IRF9.

Several IRF2-regulated genes in the RNAseq data set have anti-microbial activities. We noted significantly enhanced expression of guanylate-binding protein (GBP) family members in *Irf2*-deficient macrophages (Fig 7C)(Kim et al., 2011). Our data suggests that IRF2 inhibits expression of these anti-microbial proteins downstream of IFNAR signaling. To further investigate the GBPs, and other candidate genes identified from our RNAseq, we executed a screen searching for genes that suppressed Lm growth restriction in *Irf2*-deficient macrophages. However, we did not measure restored Lm growth in double knockout macrophages lacking any of the genes tested (Fig S5E).

The IL-21 signaling pathway was enriched in the subset of genes that are dependent on IRF9 but completely inhibited by IRF2. The role of IL-21 signaling in macrophages is not well established, however there is evidence that it can boost proinflammatory responses (Yang et al., 2023; Jian et al., 2021). The genes for both IL- 21R and the common gamma chain subunit that form the functional heterodimeric receptor are enriched in *Irf2*-deficient macrophages during Lm infection (Fig 7D). We generated *Irf2* x *Il21r* DKO CIMs and did not observe suppression of the Lm growth restriction phenotype, though this was not surprising as macrophages are not known to generate IL-21 (Fig 7E). Addition of IL-21 did not suppress Lm levels in macrophages, in fact the amount of Lm was enhanced (Fig 7E and 7F). We speculate there may be increased initial phagocytosis of the bacteria as IL-21 has been reported to enhance phagocytosis (Vallières and Girard, 2013). However, whether increased phagocytosis is the reason for enhanced Lm levels remains to be confirmed.

To replicate in a macrophage Lm must escape from the phagosome into the host cytosol (Nguyen et al., 2019). As we measure decreased Lm growth in *Irf2*-deficient macrophages as early as two hours post infection (Fig 6A and 6B), we wondered whether Lm is unable to escape the phagosome in *Irf2* KO macrophages. To test this possibility, we used a microscopic approach previously demonstrated to detect phagosomal damage induced by Lm infection (Davis et al., 2012). After late endosomes and lysosomes were loaded with fluorescein-dextran, macrophages were infected with Lm, and phagosomal damage induced by Lm allowed fluorescein-dextran to reach the cytosol. We measured less phagosomal damage induced by Lm infection in *Irf2* KO BMMs compared to WT, suggesting that that Lm escape to the cytosol is repressed in *Irf2*-deficient macrophages (Fig 7G). We wondered whether this reduced escape was due to increased renitence, an inducible membrane resistance to damage (Davis et al., 2012). However, when we assessed the phagosomal damage caused by silica microspheres we did not measure any significant differences between macrophage genotypes, indicating that the reduction in Lm-induced damage seen in *Irf2*-deficient macrophages is not due to host activities that reinforce the phagosome. One possible mechanism that could be responsible for this enhanced phagosomal killing is LC3- associated phagocytosis (LAP), in which LC3 is conjugated to phagosome membranes. LAP has been described to facilitate lysosomal fusion and enhanced microbial killing.

However, we saw no reversal in Lm restriction in macrophages lacking both *Irf2* and genes required for LAP (Fig S5E). Overall, we measured enhanced expression of anti- microbial mediators in *Irf2*-deficient macrophages. Our data suggest that one or more of these mediators kill bacteria in the phagosome prior to bacterial escape into the cytosol.

## DISCUSSION

We generated an extensive catalog of new inflammatory regulators that modulate macrophage responses to Mtb infection by screening for genes that impact TNF and iNOS induction using a genome-wide CRISPR knockout library. This dataset should be a valuable resource for those seeking to understand previously unexplored pathways that govern inflammatory responses generated by this critical cellular niche.

One of the most striking results of our screen was the correlation of IFNAR signaling components and TNF production as these pathways are often described to be antagonistic (Cantaert et al., 2010; Feng et al., 2024). However, there are also conflicting reports in which IFNAR signaling correlates with enhanced pro-inflammatory cytokine production (McNab et al., 2014; Mancuso et al., 2007). Our data provide further insight into this dichotomy as we found that type I IFN can indeed boost TNF production when there is no prior IFN signal sensed by the macrophage. In this case IFNAR signaling can mimic IFNγ signaling and induce NO production and augment the production of pro-inflammatory cytokines. We also find that in the absence of IRF2 IFNAR signaling can induce anti-bacterial mechanisms. We hypothesize that the outcome of type I IFN signaling is context-dependent and based on the temporal order and quantity of type I IFN expression relative to the engagement of other signaling pathways, especially IFNγ. We propose that the context in which IFNAR signaling can augment innate antibacterial responses is during the earliest innate immune responses, prior to the recruitment of adaptive immune cells that express IFNγ or a strong type I IFN signal that promote coherent anti-bacteria or antiviral responses, respectively, later in infection. A bacterial infection can induce NFκB signaling and, in some cases, also initiate a low-level type I IFN response. Bacteria can induce early type I IFN in a variety of ways including engagement of cytosolic or endosomal nucleic acid sensors or stimulation of TLR4 by gram negative bacteria. We propose that when a host is infected with bacteria IFNAR signaling occurs with similar kinetics as NFκB signaling and this IFNAR signal augments proinflammatory responses and antibacterial defenses.

However, a viral infection can induce high levels of type I IFN, which can reach macrophages that have not yet received an NFκB inducing signal, this alerts macrophages to initiate an anti-viral response in which IFNAR signaling does not augment proinflammatory responses. The ability to induce some proinflammatory and anti-bacterial responses in the critical early stages of a bacterial infection, prior to IFNγ production by T cells, could help the macrophage respond appropriately to diverse microbes to blunt early growth. Once the adaptive immune response has been initiated, type I IFN signaling dampens type II IFN responses. The anti-bacterial role of IFNAR signaling prior to induction of an adaptive immune response are supported by two previous studies that show a survival disadvantage in Mtb infected mice that lack both IFNAR and IFNγ signaling compared to IFNγ signaling alone; mice infected with an Mtb strain that induces high levels of type I IFN also showed increased bacterial burden in mice lacking both IFNAR and IFNγ signaling compared to IFNγ alone (Desvignes et al., 2012; Moreira-Teixeira et al., 2016).

Regulation of IRF2 levels provides another mechanism that dictates the outcome of signaling responses in a macrophage. We have found that there are conditions in which type I IFN signaling can lead to anti-bacterial responses, some of which are inhibited by IRF2. IRF2 expression could therefore be a critical factor determining when type I IFN signaling has anti-bacterial effects. Because IRF2 expression is impacted by inflammatory signaling (Cui et al., 2018), perhaps the inflammatory environment of a tissue can influence IRF2 levels and thereby impact the outcome of type I IFN signaling. IRF2 protein levels in different tissue resident macrophages may also impact how particular types of macrophages respond to IFNAR signaling.

Intriguingly, we found that multiple components of mitochondrial respiratory chain complex I is required for TNF production upon infection with Mtb. Several mitochondrial respiratory chain complex I genes were also hits in a recently published screen examining macrophage survival after Mtb infection, suggesting there may be reduced Mtb growth in these macrophages, which may result in lower production of TNF (Simwela et al., 2024). Mitochondrial respiratory chain complex I is also the target of metformin, a drug that has been demonstrated to reduce Mtb growth (Singhal et al., 2014). Alternatively, changes in the levels of mitochondrial reactive oxygen species, which can be produced by complex I, may alter the cells’ ability to produce TNF as mitochondrial reactive oxygen species have been shown to promote the production of proinflammatory cytokines (Bulua et al., 2011).

Six genes identified in our screen are members of the Spt-Ada-Gcn5 acetyltransferase (SAGA) complex, a transcriptional coactivator with multiple modules that perform independent activities. We identified *Atxn7l1, Atxn7l3,* and *Usp22* from the deubiquitylase module, *Tada2b* from the histone acetyltransferase module, *Taf6l* from the core structural module, and *Sf3b5* from the splicing module (Chen and Dent, 2021). Intriguingly, macrophages lacking these genes were enriched in TNF^+^ cells, suggesting the SAGA complex dampens TNF production. In our secondary screen we confirmed the role of the SAGA complex in TNF production using sgRNAs targeting *Atxn7l3* and measured dramatically heightened production of TNF in cells lacking *Atxn7l3* during both Mtb and Lm infection (Fig 2 and 3). Further study is needed to reveal the mechanism of SAGA-mediated inhibition of TNF production, it is possible that the SAGA complex promotes expression of a TNF repressor. Results from our screen suggest that SAGA activity may be another manner in which inflammatory cytokine production can be dynamically regulated.

In summary, our study expands our understanding of the mediators and inhibitors of inflammatory responses generated by macrophages during the early innate immune response to Mtb infection. As we know that the inflammatory environment in host tissue can dramatically impact the course of Mtb infection, insights into how inflammation is initiated and regulated could prove vital to the establishment of novel treatments. In addition to the candidates we explored, our dataset also highlights several other genes and cellular pathways that can be further investigated to determine their involvement in inflammatory responses. In addition to their potential role in Mtb infection, genes identified in this study could be involved in other inflammatory conditions including autoinflammatory disorders or cancers associated with inflammation.

## MATERIALS AND METHODS

### Mammalian cell culture

Conditionally-immortalized macrophages (CIMs) were derived as described previously (Roberts et al, 2019, Wang et al, 2006). Progenitor CIMs, prior to differentiation, were maintained in RPMI (Gibco) supplemented with 10% FBS, 2% GM-CSF supernatant produced in by a B16 murine melanoma cel line, 2mM GlutaMAX (Gibco), 1mM sodium pyruvate (Gibco), 10mM HEPES (Gibco), 40 uM β- mercaptoethanol (Sigma-Aldrich # M6250), and 2 uM β-estradiol (Sigma #E2758). Progenitor CIMs were maintained in suspension in non-TC treated tissue culture flasks at densities below 5x10^5^ cells/ml before removal of β-estradiol and differentiation. To differentiate progenitor CIMs into macrophages, cells were washed twice in DMEM + 5% FBS to fully remove β-estradiol, resuspended in macrophage media [DMEM (Gibco) supplemented with 10% FBS, 10% M-CSF supernatant produced by 3T3-MCSF cells, 2 mM GlutaMAX (Gibco), and 1 mM sodium pyruvate (Gibco)], and seeded onto non-TC treated 15 cm tissue culture plates at 3.0×10^6^ cells/plate in 25 ml of macrophage media. Differentiating CIMs were given an additional 12 ml of macrophage media on day 3 post-differentiation, and CIMs were replated for further experiments on day 7 post differentiation.

BMMs were differentiated from mouse bone marrow isolated from femurs and tibias. All mice were housed in specific-pathogen free conditions and treated using procedures described in animal care and use protocols approved by the Institutional Animal Care and Use committee of the University of California, Berkeley. *Irf2* KO mouse bone marrow was a gift from Dr. Vishva Dixit’s laboratory (Kayagaki et al., 2019), and *Lyz2cre*^+/−^*Traf3^fl/fl^* mouse bone marrow was a gift from Dr. Ping Xie’s laboratory (Lalani et al., 2015). RBCs from bone marrow cells were lysed and all remaining cells were plated in 25ml of macrophage media supplemented with 100 units/ml of penicillin and 100 μg/ml of streptomycin for 24 hours. After 24 hours, all non-adherent bone marrow cells were collected, washed in complete macrophage media to remove antibiotics, and seeded onto non-TC treated 15 cm tissue culture plates at 4.0×10^6^ cells/plate in 25 ml of media. Differentiating BMMs were given an additional 12 ml of macrophage media on day 3 post-differentiation, and were replated for further experiments on day 7 post differentiation or frozen in 5% DMSO + 95% FBS for future experiments. Frozen differentiated BMMs were thawed into 25ml complete macrophage media and harvested for future experiments 3 days post thawing.

### Genome editing of CIMs

Gene KO CIMs were generated as described in (Roberts et al., 2020) with minimal modifications. Briefly, CRISPR guide sequences targeting genes of interest were selected from the murine Brie guide library (Addgene #73633). Oligonucleotides encoding the chosen gRNAs were cloned into pLentiGuide-Puro (Addgene #52963) or lenti-gRNA hygro (Addgene #104991) and verified by sequencing using the human U6 sequencing primer. 293 T cells were co-transfected with pLentiGuide-Puro, psPAX2, and pMD2.G using Lipofectamine 3000 and Optimem according to manufacturer’s guidelines to generate lentiviral particles for transduction into Cas9-expressing CIM progenitors. For transduction, lentivirus was combined with 5.0×10^5^ CIM progenitors/well in a 6-well non-treated plate and spinfected at 1000xg for 30min at 32°C in the presence of 10 µg/ml protamine sulfate. To generate double knockout cells two different lentiviruses with differing antibiotic resistance were combined with progenitors in the same well. Two days post-transduction, 12 µg/ml puromycin and/or 250 µg/ml hygromycin was added to cells, and cells were selected for 4-7 days (until control well with no guide plasmid has completely died, hygromycin selection took longer than puromycin). Antibiotic-resistant cells were maintained as polyclonal populations and total gDNA was extracted using DNeasy Blood and Tissue kit (Qiagen). To ensure successful genome editing, genomic sites encompassing targeted guide regions were amplified by PCR using iProof polymerase (BioRad) and sequenced, and population level genome editing was established using EditCo’s ICE Analysis tool (ICE CRISPR Analysis. 2025. v3.0. EditCo Bio, www.editco.bio/crispr-analysis).

### Macrophage plating

CIMs and BMMs were harvested 7 days post differentiation using 0.05% Trypsin and plated in TC-treated black-walled plates (PhenoPlate cat # 6055300, Revvity) for microscopy experiments, clear 96-well TC-treated plates (Corning cat # 3595) for stimulation experiments, clear 24-well TC-treated plates (Corning cat # 3526) for CFU enumeration experiments, or 75cm^2^ non-TC trated flasks (Genessee) for the genome-wide screen. Macrophages were plated at 4x10^4^ cells per well in 96 well plates, 2.4x10^5^ cells per well in 24-well plates, or 14x10^6^ in 75cm^2^ flasks. For phagosome damage experiments BMMs were plated at 4x10^4^. However, when required, plating numbers were varied to ensure equivalent cell numbers on the day of infection or stimulation. For experiments with substantial numbers of knockout macrophage lines, to ensure matching cell density, control cells were plated at several different densities. This allowed for matching of knockout cells lines to control wells with equivalent numbers of cells. For Mtb infections, parallel plates were made such that on day 0 of infection one plate was fixed in 4% paraformaldehyde in PBS, washed, DAPI stained, and nuclei were counted at 10X magnification on an Opera Phenix microscope. For bacterial infections in 96-well plates cell lines were plated in random order to minimize potential effects based on well location within the plate. After plating, cells were allowed to adhere and recover for 2 or 3 days prior to infection or stimulation.

### Bacterial infections of macrophages

#### Mycobacterium tuberculosis

All Mtb experiments were performed with Mtb Erdman strain or strains derived from the Erdman strain. Mtb expressing eGFP under control of the MOP promoter was a gift from Dr. Sarah Stanley’s laboratory. Mtb expressing mCherry under control of the Mycobacterium Strong Promoter was a gift from Dr. Russell Vance’s laboratory. Low passage frozen stocks were grown to mid-log phase in 7H9 liquid media (BD) supplemented with 10% Middlebrook OADC (Sigma), 0.5% glycerol, and 0.05% Tween- 80 in roller bottles at 37°C. On the day of infection Mtb cultures were washed twice in PBS, gently centrifuged to remove large clumps, gently sonicated to disperse clumps, and resuspended in macrophage media.

For microscopy experiments macrophages were infected at an MOI of 1. To avoid disturbing macrophage monolayers full media changes were not performed during Mtb infections. Cells were infected by removing half (100μl) of the media from wells in 96 well plates, overlaying monolayers with 100μl bacterial suspensions, spinning Mtb onto macrophages at 1200 rpm for 5 min, and performing another half media change by replacing 100μl of media with fresh Mac media +/- IFNγ. Where appropriate, IFNγ was added to achieve a final concentration of 2ng/ml. On day two of infection a 75% media change was performed. Plates were fixed in 4% paraformaldehyde on day 4. After fixation, plates were washed in PBS. To stain nuclei, Dapi was added at least 30 min before imaging. Mtb infections for genome-wide screen are described in the CRISPR screen section.

### Listeria Monocytogenes

The parental strain for all Lm strains used in this study is 10403S. Strains were grown overnight, slanted at 30°C, in Bacto Brain Heart Infusion (BD #237500) supplemented with 200 μg/mL streptomycin to an OD600 of ∼1.5. Lm expressing TagRFP under control of the actA promoter was a gift from Dr. Daniel Portnoy’s Laboratory.

Lm CFU enumeration: 2.4x10^5^ BMMs or CIMs were plated (see macrophage plating section) in tissue-culture treated 24 well plates (Corning, # 3526). Two days later, BMMs or CIMs were infected at a multiplicity of infection (MOI) of 0.25 or 1.0, respectively, by incubation with macrophage media containing WT Lm for 0.5 hours. At 0.5 hours the macrophages were washed with warm 5% FBS in DMEM then incubated in fresh macrophage medium. To kill extracellular bacteria, one hour post-infection macrophage medium containing gentamicin was added to the wells to achieve a final concentration of 50 µg/mL. CFU was enumerated at 0.5 hour, 2 hour, and 5 hours post-infection. At each timepoint cells were washed with 1mL of warm PBS, then macrophages were lysed by incubating in 500µL of sterile filtered 0.05% Triton-X 100 in water for 10 minutes. Lysate was serial diluted as needed in PBS and bacteria was plated onto LB agar plates supplemented with 200µg/mL streptomycin.

Lm Microscopy, LDH, and phagosome damage experiments: BMMs or CIMs were plated (see macrophage plating section) on TC-treated black-walled plates (PhenoPlate cat # 6055300, Revvity). BMMs or CIMs were infected two days later at a multiplicity of infection (MOI) of 3.75 or 37.5, respectively, by incubation with macrophage media containing TagRFP-Lm (microscopy) or WT Lm (LDH and phagosome damage) for 0.5 hours. At 0.5 hours macrophages were washed with warm 5% FBS in DMEM then incubated in fresh macrophage medium. To kill extracellular bacteria, one hour post-infection macrophage medium containing gentamicin was added to the wells to achieve a final concentration of 50 µg/mL. For microscopy experiments plates were fixed in 4% paraformaldehyde at 5 hours post infection. After fixation plates were washed in PBS. To stain nuclei, Dapi was added at least 30 min before imaging.

Microscopy for fluorescent bacteria was performed using the Opera Phenix High-Content Screening System confocal microscope (PerkinElmer) at 40X magnification. A maximum intensity projection was generated from multiple z-stacks. Automated analysis for determination of bacterial area and cell counting was performed with PerkinElmer Harmony software packages. Cell numbers were determined based on DAPI staining of nuclei.

### Genome-wide CRISPR screens and analysis

#### Generation of genome-wide knockout library in CIMs

The mouse Brie knockout CRISPR pooled library on the lentiGuide-Puro backbone was purchased from Addgene and amplified by the UC Berkeley High Throughput Screening Facility. sgRNAs were PCR amplified and sequenced to verify sgRNA representation on the Illumina HiSeq4000, 50SR at the UC Berkeley Genomics Sequencing Laboratory.

The Brie library was transduced into CIM progenitors as described in genome editing of CIMs section with the following changes. 293T cells were transfected in 7 10cm plates with 7μg of the BRIE knockout library on lentiGuide-Puro + 4.6 μg of psPAX2 + 2.5μg pMD2.G per plate. 96 wells of ∼5x10^6^ progenitors per well were each transduced with ∼800μl lentivirus containing supernatant in a total of 2ml progenitor media. Progenitors were spinfected for 2 hours at 1000xg at 32°C in the presence of 10 µg/ml protamine sulfate. After spinfection, progenitors were immediately resuspended and all wells were combined and aliquoted into 16 total 182cm^2^ suspension culture flasks (Genessee). In total ∼470x10^6^ progenitors were transduced. One day post transduction all cells were re-combined and expanded to a concentration of 0.15x10^6^ cells/ml and put into fresh flasks. Two days post transduction all cells were combined in a sterile 10L carboy and diluted to 0.25x10^6^ cells/ml in media containing puromycin at a final concentration of 12μg/ml. Three days post puromycin addition, all cells in a test well of untransduced progenitors were dead and all library transduced progenitors were combined in a sterile 10L carboy, then centrifuged, and frozen. During selection small scale parallel flasks +/- puromycin were set up to calculate transduction efficiency. Based on transduction efficiency, the library was generated at a coverage of ∼400 progenitors/ guide. The library was frozen at >750x coverage for future experiments.

### Genome-wide CRISPR screen of macrophage differentiation

Brie library transduced progenitors were thawed and expanded for three days. Progenitors were combined, a day 0 sample of 50x10^6^ cells was collected, and 40x10^6^ (a coverage of ∼500x) progenitors were differentiated by washing twice in PBS + 2% FBS and resuspending in 20ml macrophage media in 14 15cm non-TC treated plates with 3x10^6^ cells per plate. On day 3 of differentiation 10ml fresh macrophage media was added per plate of differentiating cells. Starting on day 0 of differentiation 40x10^6^ cells were kept in progenitor media to maintain cells in a progenitor state. Progenitor cells were combined and diluted to maintain 40x10^6^ cells daily for seven days. On day 7 cells were harvested. Undifferentiated progenitors were harvested by centrifuging suspension cells and combining all cells before counting and centrifuging 50x10^6^ cells.

Differentiated macrophage plates were washed once in warm PBS and macrophages were harvested using trypsin. After harvest all cells were combined and counted, and 50x10^6^ cells were centrifuged. Cell pellets were frozen at -80C until collection of genomic DNA. In total three separate samples were collected in every screen repeat. #1) on day 0 50x10^6^ undifferentiated progenitors. #2) on day 7 50x10^6^ differentiated macrophages. #3) on day 7 50x10^6^ undifferentiated progenitors that had been passaged for 7 days. Three independent replicates of the differentiation screen were performed.

### Genome-wide CRISPR screen of TNF and iNOS induction after Mtb infection

Brie library transduced progenitors were thawed and expanded for 2-3 days prior to differentiation of 135x10^6^ progenitors. After differentiation for 7 days CIMs were harvested using trypsin and ∼500x10^6^ (∼6250x coverage) total CIMs were plated in 75cm^2^ non-TC treated flasks at 14x10^6^ macrophages per flask. Three days post plating CIMs were infected with WT Mtb at an MOI of 1.5 by addition of macrophage media containing Mtb. To establish sorting gates, one flask of CIMs was mock infected with macrophage media containing no Mtb then harvested and stained in the same manner as infected CIMs. 18 hours after infection, golgiplug (BD) was added to flasks according to manufacturer’s instructions. Six hours after addition of golgiplug, CIMs were harvested with trypsin then fixed for 20 min at 4°C at 1x10^7^ cells/ml using the cytofix/cytoperm fixation/permeabilization kit (BD #554714). Fixed cells were permeabilized by washing in perm/wash buffer (BD #554714), then blocked for 10min at room temperature using anti-CD16/32 Fc blocking antibody (clone 93, BioLegend) in perm/wash buffer. Cells were then stained for TNF (clone MP6-XT22, eBioscience # 50-112-9483) and iNOS (clone CXNFT, eBioscience # 17-5920-82) in perm/wash buffer for 30 min at room temperature. Cells were washed in perm/wash buffer then resuspended in PBS + 2% FBS + 2mM EDTA. An unsorted sample of 40x10^6^ library cells was taken. Remaining cells were then sorted into TNF+iNOS- and TNF+iNOS+ populations on a SONY SH800 sorter using uninfected library CIMs as well as single color staining controls to set sort gates. Populations were sorted at coverage of >300x for the TNF+iNOS- population and 25-75x for the TNF+iNOS+ population. After sorting, cells were again fixed in 4% paraformaldehyde for removal from the BSL3. Two independent replicates of the TNF and iNOS induction screen were performed.

### gDNA harvest

Screen sample cells were lysed overnight at 55°C at 5x10^6^ cells/ml in NK lysis buffer (50mM Tris, 50mM EDTA, 1% SDS, pH 8) + 100μg/ml proteinase K (Zymo). After lysis, RNase (Qiagen) was added for a final concentration of 50μg/ml. Samples were mixed by inverting 25 times then incubated at 37°C for 30min. After incubation, samples were cooled on ice and cold 7.5M ammonium acetate was added at a 1:4 ratio of ammonium acetate to sample, for a final concentration of 1.875M ammonium acetate. Samples were vortexed, then samples were split into 800μl aliquots in Eppendorf tubes and centrifuged at 12,000xg for 10min. Supernatants were moved into fresh Eppendorf tubes and 600μl of 100% isopropanol was added to each Eppendorf. Tubes were inverted 50 times to precipitate gDNA. Samples were then centrifuged at 20,000xg for 30min at 4°C. After centrifugation, the gDNA was visible as a small white pellet and supernatant was discarded. gDNA was washed by adding 600μl 70% ethanol per tube, inverting 10 times, and centrifuging at 20,000xg for 10min at 4°C. Ethanol was removed and gDNA was air dried for 10-30 min. gDNA was then resuspended in 10mM Tris HCL pH 8.5 + 0.1mM EDTA at 50μl per tube. Samples were incubated first at 65°C for 1 hr, then at room temperature overnight to fully resuspend gDNA. After overnight incubation all tubes for each sample were re-combined.

### PCR of sgRNA for sequencing

gDNA sample concentrations were determined by qubit. sgRNA amplification by PCR was performed with 1.5μg gDNA per 50μl reaction as previously described, using Illumina compatible primers from IDT (Doench et al., 2016). Each sample was amplified using a P7 primer with a unique barcode. All amplicons for each sample were combined and PCRs were purified using Zymo-spin V columns with reservoirs (Zymo # C1016- 25). Amplicons were then purified by agarose gel extraction using the QIAquick gel extraction kit (Qiagen # 28704). After gel extraction, amplicons were again purified and concentrated using the DNA clean & concentrator – 5 kit (Zymo #D4003). DNA was eluted in 10mM Tris-HCL pH 8.5 with no EDTA. Amplicons were sequenced at the UC Berkeley Vincent J. Coates Genomics Sequencing Laboratory on an Illumina NovaSeq 6000, 100SR.

### Screen Analysis

CRISPR sequence results were mapped and quantified using the R package MAGeCK (version 0.5.9.5). MAGeCK default mapping settings were used to automatically determine the trimming length and sgRNA length. MAGeCK’s robust rank algorithm (RRA) was then used to score genes by negative and positive enrichment with non-targeting gRNAs included as a control.

### RNAseq

Total RNA was isolated using TRIzol (Fisher) and Chloroform, and the PureLink RNA Mini Kit (12183018A, Ambion). RNA was then DNase I treated (NEB), and purified using RNA clean and concentrator columns (Zymo). PolyA mRNA was enriched and sequencied by Novogene. RNA libraries were sequenced on an Illumina NovaSeq using 150 paired end reads. The raw reads were pre-processed using HTStream (version 1.3.3) to filter out polyA tails, reads from PhiX, adapter sequences, reads that were too short (<50 bp), read ends with low quality (<Q20), and reads resulting from PCR duplication. The pre-processed reads were mapped to the Gencode M27 Mus musculus genome (GRCm39) and quantified using STAR aligner (version 2.7.9a). The R package edgeR was used to determine differentially expressed genes. Normalization was performed using trimmed mean of M-values.

### Pathway enrichment analysis

Pathway enrichment analysis was performed using Metascape (Zhou et al., 2019). Only pathways with enrichment significance of log(q-value) less than -1 were considered. For pathways with substantial gene overlap only one pathway was shown after consideration of pathway specificity as well as statistical significance. Analysis was performed on genes with a RRA significance score of less than 1x10^-3^ in the genome-wide CRISPR screen. For RNAseq differential expression was defined as at least 2 fold change and adjusted p value <0.05. Gene clusters are based on positive or negative differential expression relative to scramble.

### TLR ligand and IFN stimulations

Macrophages were seeded at 4x10^4^ cells per well in 96-well Tissue culture plates (Corning #3595) in macrophage media two days prior to stimulation. When prestimulations were performed, cytokines were added one day prior to stimulation. For cytokine analysis cells were stimulated for 20 hours with 100 ng/ml LPS (InvivoGen, # tlrl-pb5lps), 1μg/ml Pam3CSK4(Invivogen, # tlrl-pms), 10μg/ml MDP (Invivogen # tlrl-mdp), or 500 nM CpG-1668 (InvivoGen, Tlrl-1668) for BMMs or 20nM CpG-1668 for CIMs. When stimulated with cytokines macrophages were stimulated with 1000 U/ml IFNβ (pbl, #12405-1), 2ng/ml IFNγ (preprotech, #315-05), or 50 ng/ml IL-21 (preprotech, #210-21).

### Cytokine analysis

TNF ELISAs were performed using cell-free supernatants. 96-well Nunc MaxiSorp flat-bottom plate (44-2404-21, Invitrogen) were coated with anti-TNF capture antibodies (eBio 1F3F3D4 cat #14-7325-85) diluted to 3μg/ml in 0.1M sodium phosphate buffer pH 8.0 and incubated overnight at 4°C. The following day, plates were washed 3x with PBS + 0.05% Tween-20 and blocked with PBS + 1% BSA for 4 hr at room temperature. Standard measurements made with serially diluted recombinant TNF (R&D, #410-MT), as well as experimental supernatants were added to blocked plates and incubated overnight at 4°C. After 3X washes in PBS/BSA, biotin-conjugated sandwich anti-TNF (ebio MP6-XT22 and MP6-XT3 cat#13-7326-85) was added and plates were incubated for one hour. Biotin was then detected by addition of Streptavidin-HRP diluted 1:3000 (BD Biosciences). ELISA was then incubated with 50 µl of 1mg/ml OPD (Sigma) and 1μl/ml 30% hydrogen peroxide. Once developed, the reaction was stopped using 20 µl 3M HCl. Absorbance at 490 nm was read on a Tecan plate reader.

IFNβ levels, and all cytokines for secondary screening, were measured using the LEGENDplex mouse inflammation panel (Biolegend #740446) according to the manufacturer’s instructions with the following modifications. The recommended reagent and sample volumes were halved. Samples and standards were incubated with the beads overnight at 4°C. Samples and beads were stationary during incubations. Analyses were performed on a SONY SH800 or BD LSR Fortessa Analyzer.

### Griess assay

Cell-free supernatants were analyzed by Griess reaction to detect nitrite as a proxy for NO production. A solution of 0.2% napthylethylenediamine dihydrochloride was mixed 1:1 with a 2% sulfanilamide/4% phosphoric acid solution. 50 µl of this solution was mixed with 50 µl of sample supernatant and absorbance at 546 nm was measured on a Tecan plate reader. Nitrite concentrations were determined using a standard curve of sodium nitrite.

### Flow cytometry for F/480

Cells were harvested from non-TC treated plates using cold PBS + 2mM EDTA and resuspended in PBS + 2% FBS (v/v) + 2mM EDTA. Cells were blocked for 10min at 4**°**C using anti-CD16/32 Fc blocking antibody (clone 93, BioLegend) and stained for 30 min at 4**°**C with antibodies against F4/80 (clone BM8, Tonbo Biosciences). After staining cells were washed and resuspended in PBS + 2% FBS (v/v) + 2mM EDTA with DAPI (1:1000) to stain dead cells. Cells were analyzed on an LSR Fortessa (BD Biosciences), and data was analyzed with FlowJo.

### LDH

After 5 hours of Lm infection, macrophage lysis was assessed using the CyQuant LDH Cytotoxicity Assay (Invitrogen, #C20300) according to the manufacturer’s instructions.

### Phagosome damage analysis

BMMs were loaded with fluorescein-dextran with an average molecular weight of 3kD (Invitrogen #D3305) for 18 hours. Cells were then washed twice in DMEM + 10% FBS and incubated in macrophage media without fluorescein-dextran for 2 hours. BMMs were infected with WT Lm or administered acid washed silica microspheres for 2 hours.

Silica microspheres (microspheres – nanospheres, cat # 140218-10) were acid washed by overnight incubation in 1M HCl, then washed 3 times in PBS before resuspension in macrophage media. After 2 hours, macrophages were put into warm Hanks buffered saline solution and fluorescence was measured after excitation at 425nm and 488nm. Microscopy was performed using the Opera Phenix High-Content Screening System confocal microscope (PerkinElmer) at 60X magnification. Automated analysis for pH determination and cell counting was performed with PerkinElmer Harmony software packages. A maximum intensity projection was generated from multiple z-stacks and as fluorescein fluorescence after excitation at 425nm is largely pH insensitive, while fluorescence after excitation at 488nm is pH sensitive, the ratio of fluorescence intensity at 488/425 at each location in the cell was used to determine pH in that location. A pH curve was established in BMMs using the Invitrogen Intracellular pH Calibration Buffer kit (cat # P35379). Fluorescein-dextran was determined to be in the cytosol when the pH was above 5.5 based on the 488/425 ratio determined from the pH calibration curve with only areas >700 pixels in area selected to limit background (Davis et al., 2012). Cell numbers were determined based on 425nm excitation image.

### Statistics

Statistics were performed using GraphPad Prism software. Data are reported as the mean +/- SD. Statistical tests used are noted in figure legends.

## Data availability

All CRISPR knockout screen and RNAseq data generated in this work will be available through the NCBI Gene Expression Omnibus upon publication.

## ACKNOWLEDGMENTS

We thank Vishva Dixit for providing *Irf2* KO mouse bone marrow, and Ping Xie for providing *Lyz2cre*^+/−^*Traf3^fl/fl^* mouse bone marrow. We thank Daniel A Portnoy for the TagRFP expressing Lm strain as well as valuable advice. We thank members of the Cox, Stanley, and Vance labs for constructive discussions. This work was supported by NIH Grants U19AI162583 (J.S.C.) and U19AI135990 (J.S.C.), and funding from Open Philanthropy (A.R). Allison Roberts was an Open Philanthropy Fellow of the Life Sciences Research Foundation. J.S.C. is on the scientific advisory board of Xbiotix Therapeutics. The authors have no additional financial interests.

---

**Figure S1:**
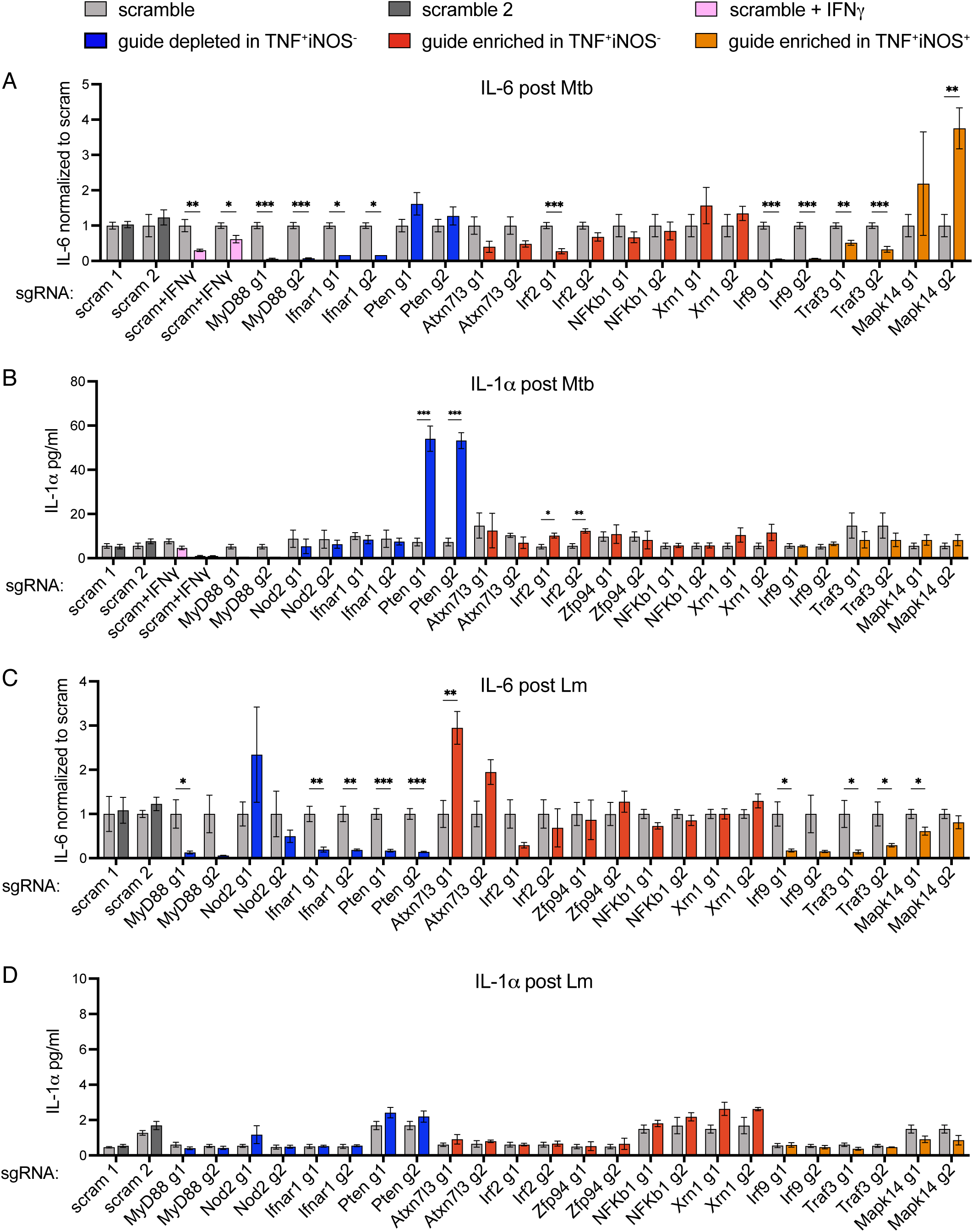
Arrayed secondary screen identifies regulators of cytokine induction upon *Mycobacterium tuberculosis and Listeria monocytogenes* infection of macrophages. CIMs lacking candidate genes were generated using two independent sgRNA for each candidate. (A and B) Two days post infection with Mtb, supernatants were measured for levels of (A) IL-6 or (B) IL-1α. (C and D) Five hours post infection with Lm, supernatants were measured for levels of (C) IL-6 or (D) IL-1α. IL-6 levels were normalized to levels generated by control (scramble gRNA) macrophages. Graphs show compiled data from four experiments, each examining different guides and the corresponding controls. Data for each guide show the mean +/- SD from one experiment with four replicates. A second independent experiment confirmed the results for *Ifnar1, Atxn7l3, Irf2, and Traf3*. Data were statistically analyzed using t-test with Holm-Sidak’s multiple comparisons test. *P < 0.05, **P < 0.01, ***P < 0.001, **** P < 0.0001.

**Figure S2:**
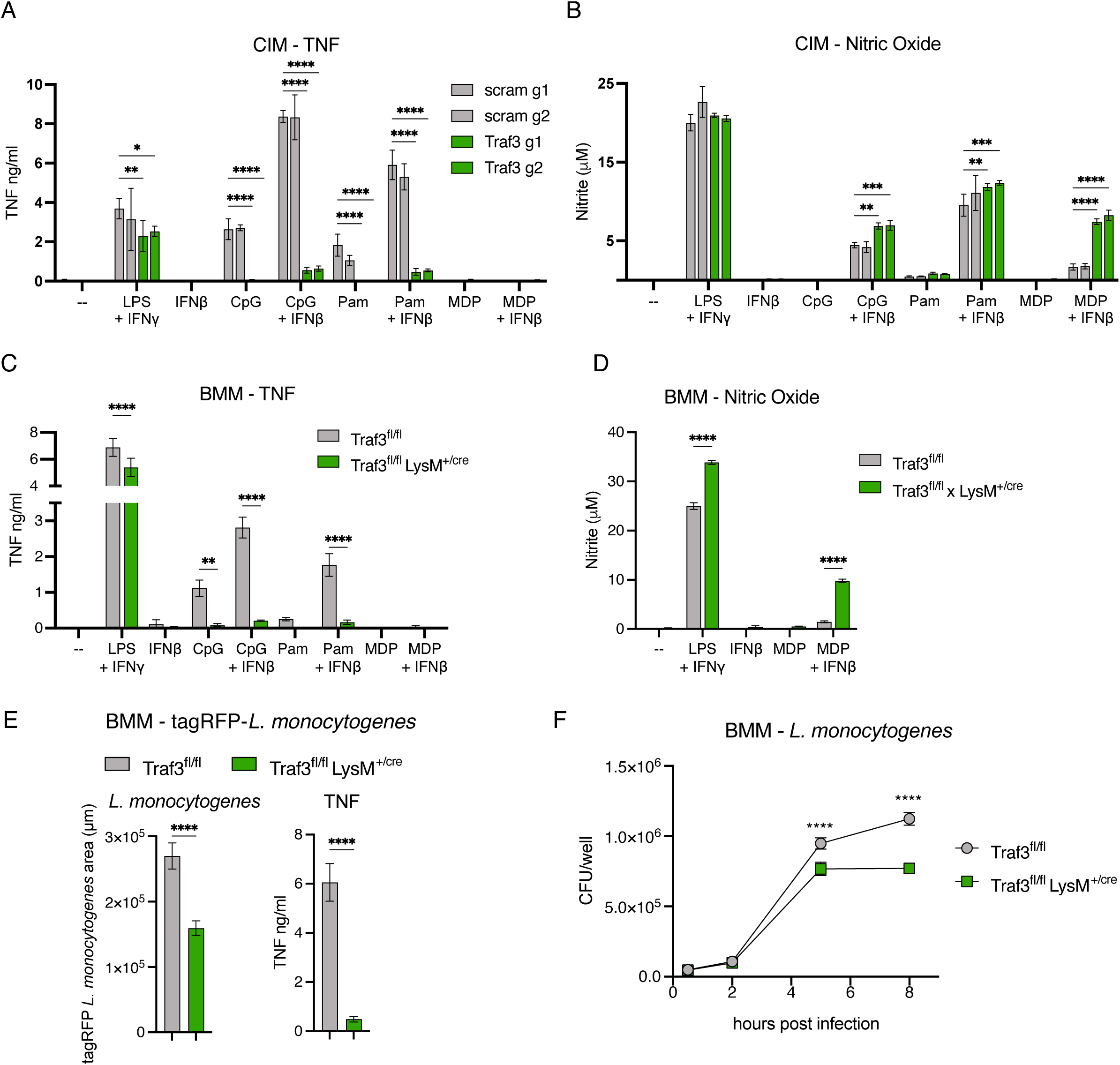
*Traf3* deficient macrophages generate reduced TNF in response to PRR stimulation, and modestly restrict *Listeria monocytogenes* growth. (A) TNF and (B) nitric oxide production by CIMs, transduced with guides for *Traf3* or scramble controls, stimulated with Pam3CSK4, CpG, or MDP +/- IFNβ, or the positive control LPS + IFNγ. Data are representative of at least two independent experiments. (C) TNF and (D) nitric oxide production by control and *Traf3* KO BMMs stimulated with Pam3CSK4, CpG, or MDP +/- IFNβ, or the positive control LPS + IFNγ. Data are representative of at least two independent experiments. (E) Control and *Traf3* KO BMMs were infected with TagRFP-Lm. Five hours post infection Lm growth in macrophages was analyzed by quantifying TagRFP area and cell supernatants were analyzed for levels of TNF. Data are representative of at least two independent experiments. (F) Control and *Traf3* KO BMMs were infected with WT Lm. Intracellular CFU were enumerated at indicated time points post infection. Data are representative of three independent experiments. All data are presented as mean +/- SD and were statistically analyzed using two-way ANOVA with Sidak’s multiple comparisons test or t-test (E). *P < 0.05, **P < 0.01, ***P < 0.001, **** P < 0.0001.

**Figure S3:**
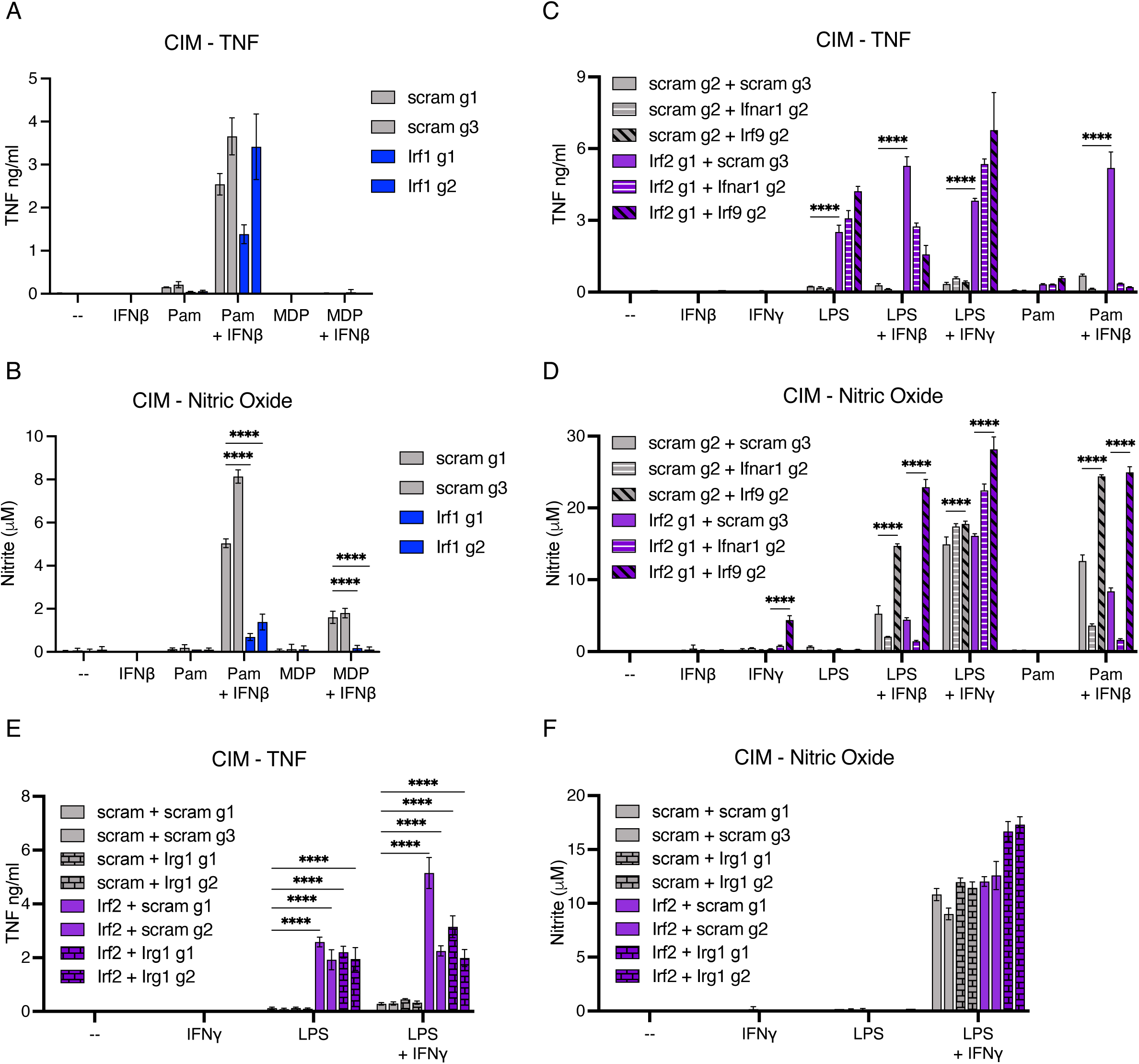
IRF2 inhibits TNF production in response to PRR stimulation. (A) TNF and (B) nitric oxide production by CIMs, transduced with guides for *Irf1* or scramble controls, stimulated with Pam3CSK4 or MDP +/- IFNβ. Data are representative of three independent experiments. (C) TNF and (D) nitric oxide production by double knockout CIMs (transduced with guides for *Irf2* or scramble and *Irf9, Ifnar1*, or scramble) stimulated with LPS or Pam3CSK4 +/- IFNβ or IFNγ. Double knockout CIMs were generated using different guides from those of Fig4C and D. Data are representative of at least three independent experiments. (E) TNF and (F) nitric oxide production by double knockout CIMs (transduced with guides for *Irf2* or scramble and *Irg1* or scramble) stimulated with LPS +/- IFNγ. Data are representative of two independent experiments. All data are presented as mean +/- SD and were statistically analyzed using two-way ANOVA with Sidak’s multiple comparisons test. *P < 0.05, **P < 0.01, ***P < 0.001, **** P < 0.0001.

**Figure S4:**
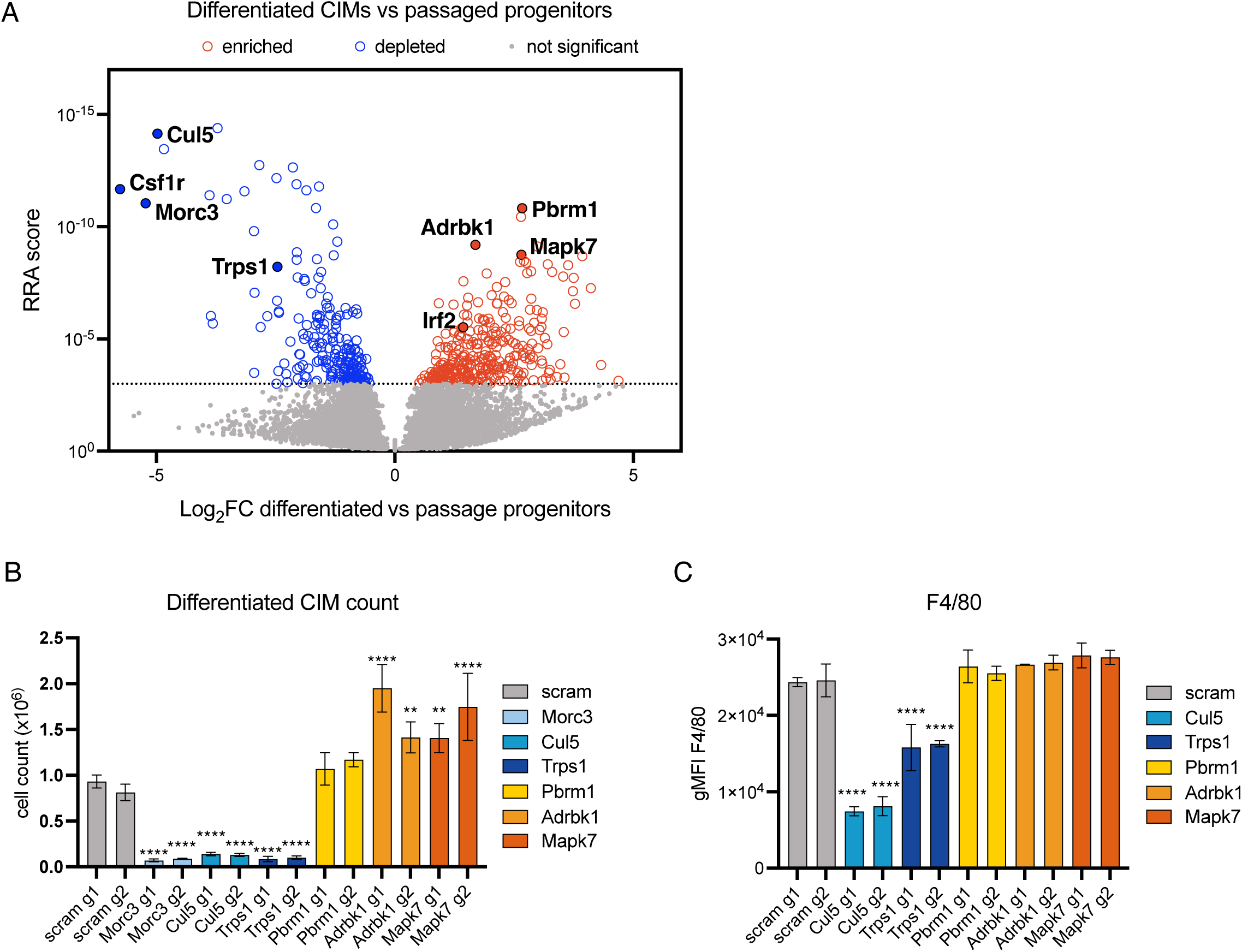
Genome-wide CRISPR knockout screen examining macrophage differentiation and proliferation. (A) The CIM progenitor knockout library was split into two groups, group 1 was differentiated into macrophages, while group 2 was passaged in the presence of estradiol to prevent macrophage differentiation. After seven days the passaged progenitors were compared to differentiated cells. Shown are results of genes depleted or enriched in differentiate CIMs compared to progenitors. Data are the combined results of three independent experiments. (B and C) CIM progenitors lacking candidate genes were generated using two independent sgRNA for each candidate, and cells were differentiated into macrophages. After seven days all suspension and adherent cells were harvested and (B) cell numbers were quantified. (C) Collected cells were stained with antibodies for F4/80 and analyzed by flow cytometry. Data are presented as mean +/- SD and were statistically analyzed using one-way ANOVA with Sidak’s multiple comparisons test. *P < 0.05, **P < 0.01, ***P < 0.001, **** P < 0.0001.

**Figure S5:**
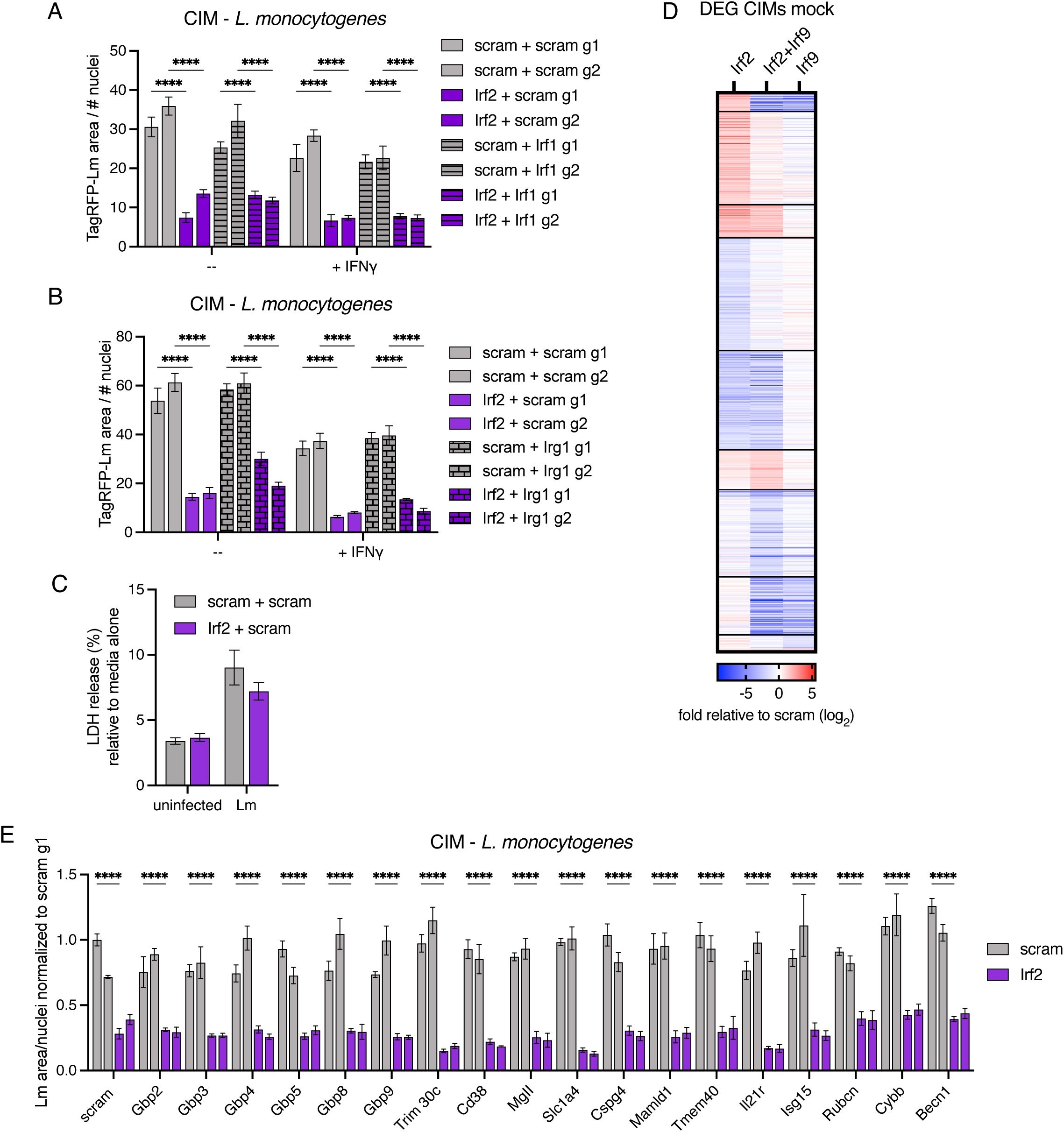
IRF2 regulates expression of antimicrobial mediators. (A and B) Double knockout CIMs (transduced with guides for *Irf2* or scramble and guides for (A) *Irf1* or (B) *Irg1* or scramble) were prestimulated +/- IFNγ before infection with TagRFP-Lm. Five hours post infection Lm growth in CIMs was analyzed by quantifying TagRFP area. Data are representative of at least two independent experiments. (C) CIMs, transduced with guides for *Irf2* or scramble, were infected with WT Lm. Five hours post infection macrophage lysis was quantified by LDH release relative to media alone. Data are representative of two experiments. (D) Average log2 fold change for knockout macrophages relative to control of genes with differential expression in at least one knockout macrophage line during mock infection. Genes are clustered based on fold change in different knockout macrophage lines. Data are from three independent experiments. (E) Double knockout CIMs (transduced with guides for *Irf2* or scramble as well as candidate genes or scramble) were infected with TagRFP-Lm. Five hours post infection Lm growth in macrophages was analyzed. Graphs show compiled data from four experiments, each examining different guides and the corresponding controls. Data from separate experiments were normalized to a scramble control CIM. All data are presented as mean +/- SD and were statistically analyzed using two-way ANOVA with Sidak’s multiple comparisons test. *P < 0.05, **P < 0.01, ***P < 0.001, **** P < 0.0001.

## REFERENCES

Barber, D.L., K.D. Mayer-Barber, C.G. Feng, A.H. Sharpe, and A. Sher. 2011. CD4 T Cells Promote Rather than Control Tuberculosis in the Absence of PD-1–Mediated Inhibition. The Journal of Immunology. 186:1598–1607. doi:10.4049/jimmunol.1003304.

Boneca, I.G., O. Dussurget, D. Cabanes, M.-A. Nahori, S. Sousa, M. Lecuit, E. Psylinakis, V. Bouriotis, J.-P. Hugot, M. Giovannini, A. Coyle, J. Bertin, A. Namane, J.-C. Rousselle, N. Cayet, M.-C. Prévost, V. Balloy, M. Chignard, D.J. Philpott, P. Cossart, and S.E. Girardin. 2007. A critical role for peptidoglycan N-deacetylation in Listeria evasion from the host innate immune system. Proceedings of the National Academy of Sciences. 104:997–1002. doi:10.1073/pnas.0609672104.

Braverman, J., and S.A. Stanley. 2017. Nitric Oxide Modulates Macrophage Responses to Mycobacterium tuberculosis Infection through Activation of HIF-1α and Repression of NF-κB. The Journal of Immunology. 199:1805–1816. doi:10.4049/jimmunol.1700515.

Bulua, A.C., A. Simon, R. Maddipati, M. Pelletier, H. Park, K.-Y. Kim, M.N. Sack, D.L. Kastner, and R.M. Siegel. 2011. Mitochondrial reactive oxygen species promote production of proinflammatory cytokines and are elevated in TNFR1-associated periodic syndrome (TRAPS). J Exp Med. 208:519–533. doi:10.1084/jem.20102049.

Cantaert, T., D. Baeten, P.P. Tak, and L.G. van Baarsen. 2010. Type I IFN and TNFα cross-regulation in immune-mediated inflammatory disease: basic concepts and clinical relevance. Arthritis Research & Therapy. 12:219. doi:10.1186/ar3150.

Carneiro, F.R.G., A. Lepelley, J.J. Seeley, M.S. Hayden, and S. Ghosh. 2018. An Essential Role for ECSIT in Mitochondrial Complex I Assembly and Mitophagy in Macrophages. Cell Reports. 22:2654–2666. doi:10.1016/j.celrep.2018.02.051.

Casey, A.M., D.G. Ryan, H.A. Prag, S.R. Chowdhury, E. Marques, K. Turner, A.V. Gruszczyk, M. Yang, D.M. Wolf, J. Lj. Miljkovic, J. Valadares, P.F. Chinnery, R.C. Hartley, C. Frezza, J. Prudent, and M.P. Murphy. 2025. Pro-inflammatory macrophages produce mitochondria-derived superoxide by reverse electron transport at complex I that regulates IL-1β release during NLRP3 inflammasome activation. Nat Metab. 7:493–507. doi:10.1038/s42255-025-01224-x.

Cavalli, G., S. Colafrancesco, G. Emmi, M. Imazio, G. Lopalco, M.C. Maggio, J. Sota, and C.A. Dinarello. 2021. Interleukin 1α: a comprehensive review on the role of IL-1α in the pathogenesis and treatment of autoimmune and inflammatory diseases. Autoimmunity Reviews. 20:102763. doi:10.1016/j.autrev.2021.102763.

Chan, J., Y. Xing, R.S. Magliozzo, and B.R. Bloom. 1992. Killing of virulent Mycobacterium tuberculosis by reactive nitrogen intermediates produced by activated murine macrophages. Journal of Experimental Medicine. 175:1111–1122. doi:10.1084/jem.175.4.1111.

Chandra, P., S.J. Grigsby, and J.A. Philips. 2022. Immune evasion and provocation by Mycobacterium tuberculosis. Nat Rev Microbiol. 20:750–766. doi:10.1038/s41579-022-00763-4.

Chen, Y.-J.C., and S.Y.R. Dent. 2021. Conservation and diversity of the eukaryotic SAGA coactivator complex across kingdoms. Epigenetics & Chromatin. 14:26. doi:10.1186/s13072-021-00402-x.

Coulombe, F., M. Divangahi, F. Veyrier, L. de Léséleuc, J.L. Gleason, Y. Yang, M.A. Kelliher, A.K. Pandey, C.M. Sassetti, M.B. Reed, and M.A. Behr. 2009. Increased NOD2- mediated recognition of N-glycolyl muramyl dipeptide. Journal of Experimental Medicine. 206:1709–1716. doi:10.1084/jem.20081779.

Cui, H., S. Banerjee, S. Guo, N. Xie, and G. Liu. 2018. Interferon regulatory factor 2 inhibits expression of glycolytic genes and lipopolysaccharide induced pro-inflammatory responses in macrophages. J Immunol. 200:3218–3230. doi:10.4049/jimmunol.1701571.

Davis, M.J., B. Gregorka, J.E. Gestwicki, and J.A. Swanson. 2012. Inducible Renitence limits Listeria monocytogenes Escape from Vacuoles in Macrophages. J Immunol. 189:4488– 4495. doi:10.4049/jimmunol.1103158.

Desvignes, L., A.J. Wolf, and J.D. Ernst. 2012. Dynamic Roles of Type I and Type II IFNs in Early Infection with Mycobacterium tuberculosis. The Journal of Immunology. 188:6205– 6215. doi:10.4049/jimmunol.1200255.

Doench, J.G., N. Fusi, M. Sullender, M. Hegde, E.W. Vaimberg, K.F. Donovan, I. Smith, Z. Tothova, C. Wilen, R. Orchard, H.W. Virgin, J. Listgarten, and D.E. Root. 2016. Optimized sgRNA design to maximize activity and minimize off-target effects of CRISPR-Cas9. Nat Biotechnol. 34:184–191. doi:10.1038/nbt.3437.

Dorhoi, A., C. Desel, V. Yeremeev, L. Pradl, V. Brinkmann, H.-J. Mollenkopf, K. Hanke, O. Gross, J. Ruland, and S.H.E. Kaufmann. 2010. The adaptor molecule CARD9 is essential for tuberculosis control. J Exp Med. 207:777–792. doi:10.1084/jem.20090067.

Drennan, M.B., D. Nicolle, V.J.F. Quesniaux, M. Jacobs, N. Allie, J. Mpagi, C. Frémond, H. Wagner, C. Kirschning, and B. Ryffel. 2004. Toll-Like Receptor 2-Deficient Mice Succumb to Mycobacterium tuberculosis Infection. Am J Pathol. 164:49–57.

Feng, J., Y. Liu, J. Kim, F. Ahangari, N. Kaminski, W.G. Bain, Z. Jie, C.S. Dela Cruz, and L. Sharma. 2024. Anti-inflammatory roles of type I interferon signaling in the lung. American Journal of Physiology-Lung Cellular and Molecular Physiology. 326:L551– L561. doi:10.1152/ajplung.00353.2023.

Flynn, J.L., M.M. Goldstein, J. Chan, K.J. Triebold, K. Pfeffer, C.J. Lowenstein, R. Schrelber, T.W. Mak, and B.R. Bloom. 1995. Tumor necrosis factor-α is required in the protective immune response against mycobacterium tuberculosis in mice. Immunity. 2:561–572. doi:10.1016/1074-7613(95)90001-2.

Gandotra, S., S. Jang, P.J. Murray, P. Salgame, and S. Ehrt. 2007. Nucleotide-Binding Oligomerization Domain Protein 2-Deficient Mice Control Infection with Mycobacterium tuberculosis. Infect Immun. 75:5127–5134. doi:10.1128/IAI.00458-07.

Global Tuberculosis Report 2024. 2024. 1st ed. World Health Organization, Geneva. 1 pp.

Gothe, F., J.S. Spegarova, C.F. Hatton, H. Griffin, T. Sargent, S.A. Cowley, W. James, A. Roppelt, A. Shcherbina, F. Hauck, H.T. Reyburn, C.J.A. Duncan, and S. Hambleton. 2022. Aberrant inflammatory responses to type I interferon in STAT2 or IRF9 deficiency. Journal of Allergy and Clinical Immunology. 150:955–964.e16. doi:10.1016/j.jaci.2022.01.026.

Häcker, H., V. Redecke, B. Blagoev, I. Kratchmarova, L.-C. Hsu, G.G. Wang, M.P. Kamps, E. Raz, H. Wagner, G. Häcker, M. Mann, and M. Karin. 2006. Specificity in Toll-like receptor signalling through distinct effector functions of TRAF3 and TRAF6. Nature. 439:204–207. doi:10.1038/nature04369.

Harris, J., and J. Keane. 2010. How tumour necrosis factor blockers interfere with tuberculosis immunity. Clinical and Experimental Immunology. 161:1–9. doi:10.1111/j.1365-2249.2010.04146.x.

Ji, D.X., K.C. Witt, D.I. Kotov, S.R. Margolis, A. Louie, V. Chevée, K.J. Chen, M.M. Gaidt, H.S. Dhaliwal, A.Y. Lee, S.L. Nishimura, D.S. Zamboni, I. Kramnik, D.A. Portnoy, K.H. Darwin, and R.E. Vance. 2021. Role of the transcriptional regulator SP140 in resistance to bacterial infections via repression of type I interferons. eLife. 10:e67290. doi:10.7554/eLife.67290.

Ji, L., T. Li, H. Chen, Y. Yang, E. Lu, J. Liu, W. Qiao, and H. Chen. 2023. The crucial regulatory role of type I interferon in inflammatory diseases. Cell Biosci. 13:230. doi:10.1186/s13578-023-01188-z.

Jian, L., C. Li, L. Sun, Z. Ma, X. Wang, R. Yu, X. Liu, and J. Zhao. 2021. IL-21 regulates macrophage activation in human monocytic THP-1-derived macrophages. Rheumatology & Autoimmunity. 1:18–29. doi:10.1002/rai2.12000.

Kamijo, R., H. Harada, T. Matsuyama, M. Bosland, J. Gerecitano, D. Shapiro, J. Le, S.I. Koh, T. Kimura, S.J. Green, T.W. Mak, T. Taniguchi, and J. Vilček. 1994. Requirement for Transcription Factor IRF-1 in NO Synthase Induction in Macrophages. Science. 263:1612–1615. doi:10.1126/science.7510419.

Kayagaki, N., B.L. Lee, I.B. Stowe, O.S. Kornfeld, K. O’Rourke, K.M. Mirrashidi, B. Haley, C. Watanabe, M. Roose-Girma, Z. Modrusan, S. Kummerfeld, R. Reja, Y. Zhang, V. Cho, T.D. Andrews, L.X. Morris, C.C. Goodnow, E.M. Bertram, and V.M. Dixit. 2019. IRF2 transcriptionally induces GSDMD expression for pyroptosis. Science Signaling. 12:eaax4917. doi:10.1126/scisignal.aax4917.

Keane, J., S. Gershon, R.P. Wise, E. Mirabile-Levens, J. Kasznica, W.D. Schwieterman, J.N. Siegel, and M.M. Braun. 2001. Tuberculosis Associated with Infliximab, a Tumor Necrosis Factor α–Neutralizing Agent. New England Journal of Medicine. 345:1098– 1104. doi:10.1056/NEJMoa011110.

Kim, B.-H., A.R. Shenoy, P. Kumar, R. Das, S. Tiwari, and J.D. MacMicking. 2011. A Family of IFN-γ–Inducible 65-kD GTPases Protects Against Bacterial Infection. Science. 332:717– 721. doi:10.1126/science.1201711.

Kobayashi, K.S., M. Chamaillard, Y. Ogura, O. Henegariu, N. Inohara, G. Nuñez, and R.A. Flavell. 2005. Nod2-Dependent Regulation of Innate and Adaptive Immunity in the Intestinal Tract. Science. 307:731–734. doi:10.1126/science.1104911.

Lalani, A.I., C.R. Moore, C. Luo, B.Z. Kreider, Y. Liu, H.C. Morse III, and P. Xie. 2015. Myeloid Cell TRAF3 Regulates Immune Responses and Inhibits Inflammation and Tumor Development in Mice. The Journal of Immunology. 194:334–348. doi:10.4049/jimmunol.1401548.

Lin, M., X. Ji, Y. Lv, D. Cui, and J. Xie. 2023. The Roles of TRAF3 in Immune Responses. Disease Markers. 2023:7787803. doi:10.1155/2023/7787803.

Loo, C.-S., J. Gatchalian, Y. Liang, M. Leblanc, M. Xie, J. Ho, B. Venkatraghavan, D.C. Hargreaves, and Y. Zheng. 2020. A Genome-wide CRISPR Screen Reveals a Role for the Non-canonical Nucleosome-Remodeling BAF Complex in Foxp3 Expression and Regulatory T Cell Function. Immunity. 53:143–157.e8. doi:10.1016/j.immuni.2020.06.011.

MacDonald, K.P.A., J.S. Palmer, S. Cronau, E. Seppanen, S. Olver, N.C. Raffelt, R. Kuns, A.R. Pettit, A. Clouston, B. Wainwright, D. Branstetter, J. Smith, R.J. Paxton, D.P. Cerretti, L. Bonham, G.R. Hill, and D.A. Hume. 2010. An antibody against the colony-stimulating factor 1 receptor depletes the resident subset of monocytes and tissue- and tumor-associated macrophages but does not inhibit inflammation. Blood. 116:3955–3963. doi:10.1182/blood-2010-02-266296.

Mancuso, G., A. Midiri, C. Biondo, C. Beninati, S. Zummo, R. Galbo, F. Tomasello, M. Gambuzza, G. Macrì, A. Ruggeri, T. Leanderson, and G. Teti. 2007. Type I IFN Signaling Is Crucial for Host Resistance against Different Species of Pathogenic Bacteria1. J Immunol. 178:3126–3133. doi:10.4049/jimmunol.178.5.3126.

Matsuyama, T., T. Kimura, M. Kitagawa, K. Pfeffer, T. Kawakami, N. Watanabe, T.M. Kündig, R. Amakawa, K. Kishihara, A. Wakeham, J. Potter, C.L. Furlonger, A. Narendran, H. Suzuki, P.S. Ohashi, C.J. Paige, T. Taniguchi, and T.W. Mak. 1993. Targeted disruption of IRF-1 or IRF-2 results in abnormal type I IFN gene induction and aberrant lymphocyte development. Cell. 75:83–97. doi:10.1016/S0092-8674(05)80086-8.

Mayer-Barber, K.D., D.L. Barber, K. Shenderov, S.D. White, M.S. Wilson, A. Cheever, D. Kugler, S. Hieny, P. Caspar, G. Núñez, D. Schlueter, R.A. Flavell, F.S. Sutterwala, and A. Sher. 2010. Cutting Edge: Caspase-1 Independent IL-1β Production Is Critical for Host Resistance to Mycobacterium tuberculosis and Does Not Require TLR Signaling In Vivo. The Journal of Immunology. 184:3326–3330. doi:10.4049/jimmunol.0904189.

McNab, F.W., J. Ewbank, A. Howes, L. Moreira-Teixeira, A. Martirosyan, N. Ghilardi, M. Saraiva, and A. O’Garra. 2014. Type I IFN Induces IL-10 Production in an IL-27– Independent Manner and Blocks Responsiveness to IFN-γ for Production of IL-12 and Bacterial Killing in Mycobacterium tuberculosis–Infected Macrophages. The Journal of Immunology. 193:3600–3612. doi:10.4049/jimmunol.1401088.

Mills, E.L., B. Kelly, A. Logan, A.S.H. Costa, M. Varma, C.E. Bryant, P. Tourlomousis, J.H.M. Däbritz, E. Gottlieb, I. Latorre, S.C. Corr, G. McManus, D. Ryan, H.T. Jacobs, M. Szibor, R.J. Xavier, T. Braun, C. Frezza, M.P. Murphy, and L.A. O’Neill. 2016. Succinate Dehydrogenase Supports Metabolic Repurposing of Mitochondria to Drive Inflammatory Macrophages. Cell. 167:457–470.e13. doi:10.1016/j.cell.2016.08.064.

Mishra, B.B., R.R. Lovewell, A.J. Olive, G. Zhang, W. Wang, E. Eugenin, C.M. Smith, J.Y. Phuah, J.E. Long, M.L. Dubuke, S.G. Palace, J.D. Goguen, R.E. Baker, S. Nambi, R. Mishra, M.G. Booty, C.E. Baer, S.A. Shaffer, V. Dartois, B.A. McCormick, X. Chen, and C.M. Sassetti. 2017. Nitric oxide prevents a pathogen-permissive granulocytic inflammation during tuberculosis. Nat Microbiol. 2:1–11. doi:10.1038/nmicrobiol.2017.72.

Moreira-Teixeira, L., J. Sousa, F.W. McNab, E. Torrado, F. Cardoso, H. Machado, F. Castro, V. Cardoso, J. Gaifem, X. Wu, R. Appelberg, A.G. Castro, A. O’Garra, and M. Saraiva. 2016. Type I IFN Inhibits Alternative Macrophage Activation during Mycobacterium tuberculosis Infection and Leads to Enhanced Protection in the Absence of IFN-γ Signaling. The Journal of Immunology. 197:4714–4726. doi:10.4049/jimmunol.1600584.

Nair, S., J.P. Huynh, V. Lampropoulou, E. Loginicheva, E. Esaulova, A.P. Gounder, A.C.M. Boon, E.A. Schwarzkopf, T.R. Bradstreet, B.T. Edelson, M.N. Artyomov, C.L. Stallings, and M.S. Diamond. 2018. Irg1 expression in myeloid cells prevents immunopathology during M. tuberculosis infection. J Exp Med. 215:1035–1045. doi:10.1084/jem.20180118.

Nguyen, B.N., B.N. Peterson, and D.A. Portnoy. 2019. Listeriolysin O: a phagosome-specific cytolysin revisited. Cell Microbiol. 21:e12988. doi:10.1111/cmi.12988.

Oganesyan, G., S.K. Saha, B. Guo, J.Q. He, A. Shahangian, B. Zarnegar, A. Perry, and G. Cheng. 2006. Critical role of TRAF3 in the Toll-like receptor-dependent and - independent antiviral response. Nature. 439:208–211. doi:10.1038/nature04374.

Orozco, S.L., S.P. Canny, and J.A. Hamerman. 2021. Signals governing monocyte differentiation during inflammation. Current Opinion in Immunology. 73:16–24. doi:10.1016/j.coi.2021.07.007.

Parrington, J., N.C. Rogers, D.R. Gewert, R. Pine, S.A. Veals, D.E. Levy, G.R. Stark, and I.M. Kerr. 1993. The interferon-stimulable response elements of two human genes detect overlapping sets of transcription factors. European Journal of Biochemistry. 214:617– 626. doi:10.1111/j.1432-1033.1993.tb17961.x.

Qin, R.-X., X.-Y. Ma, Z.-Y. Han, S.-Y. Ma, Z. Shen, Z.-H. Lu, Y. Sun, and W.-L. Yu. 2024. IRF2 Affects LPS- and IFN-γ-Induced Pro-Inflammatory Responses, Cell Viability, Migration and Apoptosis of Macrophages by Regulating IRG1. *J Inflamm Res*. 17:9651–9664. doi:10.2147/JIR.S490655.

Raymond, J.B., S. Mahapatra, D.C. Crick, and M.S. Pavelka. 2005. Identification of the *namH* Gene, Encoding the Hydroxylase Responsible for the *N*-Glycolylation of the Mycobacterial Peptidoglycan*. Journal of Biological Chemistry. 280:326–333. doi:10.1074/jbc.M411006200.

Roberts, A.W., L.M. Popov, and J.S. Cox. 2020. Gene knockout and differentiation of Cas9+ CIMs. bio-protocol.

Roberts, A.W., L.M. Popov, G. Mitchell, K.L. Ching, D.J. Licht, G. Golovkine, G.M. Barton, and J.S. Cox. 2019. Cas9+ conditionally-immortalized macrophages as a tool for bacterial pathogenesis and beyond. eLife. 8:e45957. doi:10.7554/eLife.45957.

Segueni, N., S. Benmerzoug, S. Rose, A. Gauthier, M.-L. Bourigault, F. Reverchon, A. Philippeau, F. Erard, M. Le Bert, H. Bouscayrol, T. Wachter, I. Garcia, G. Kollias, M. Jacobs, B. Ryffel, and V.F.J. Quesniaux. 2016. Innate myeloid cell TNFR1 mediates first line defence against primary Mycobacterium tuberculosis infection. Scientific Reports. 6:1–14. doi:10.1038/srep22454.

Simmons, J.D., C.M. Stein, C. Seshadri, M. Campo, G. Alter, S. Fortune, E. Schurr, R.S. Wallis, G. Churchyard, H. Mayanja-Kizza, W.H. Boom, and T.R. Hawn. 2018. Immunological mechanisms of human resistance to persistent Mycobacterium tuberculosis infection. Nat Rev Immunol. 18:575–589. doi:10.1038/s41577-018-0025-3.

Simwela, N.V., L. Johnston, P.P. Bitar, E. Jaecklein, C. Altier, C.M. Sassetti, and D.G. Russell. 2024. Genome-wide screen of Mycobacterium tuberculosis-infected macrophages revealed GID/CTLH complex-mediated modulation of bacterial growth. Nat Commun. 15:9322. doi:10.1038/s41467-024-53637-z.

Singhal, A., L. Jie, P. Kumar, G.S. Hong, M.K.-S. Leow, B. Paleja, L. Tsenova, N. Kurepina, J. Chen, F. Zolezzi, B. Kreiswirth, M. Poidinger, C. Chee, G. Kaplan, Y.T. Wang, and G. De Libero. 2014. Metformin as adjunct antituberculosis therapy. Science Translational Medicine. 6:263ra159–263ra159. doi:10.1126/scitranslmed.3009885.

Stanley, S.A., J.E. Johndrow, P. Manzanillo, and J.S. Cox. 2007. The Type I IFN Response to Infection with Mycobacterium tuberculosis Requires ESX-1-Mediated Secretion and Contributes to Pathogenesis1. The Journal of Immunology. 178:3143–3152. doi:10.4049/jimmunol.178.5.3143.

Sun, L., Z. Jiang, V.A. Acosta-Rodriguez, M. Berger, X. Du, J.H. Choi, J. Wang, K. Wang, G.K. Kilaru, J.A. Mohawk, J. Quan, L. Scott, S. Hildebrand, X. Li, M. Tang, X. Zhan, A.R. Murray, D. La Vine, E.M.Y. Moresco, J.S. Takahashi, and B. Beutler. 2017. HCFC2 is needed for IRF1- and IRF2-dependent Tlr3 transcription and for survival during viral infections. Journal of Experimental Medicine. 214:3263–3277. doi:10.1084/jem.20161630.

Vallières, F., and D. Girard. 2013. IL-21 Enhances Phagocytosis in Mononuclear Phagocyte Cells: Identification of Spleen Tyrosine Kinase as a Novel Molecular Target of IL-21. The Journal of Immunology. 190:2904–2912. doi:10.4049/jimmunol.1201941.

Vanneste, D., Q. Bai, S. Hasan, W. Peng, D. Pirottin, J. Schyns, P. Maréchal, C. Ruscitti, M. Meunier, Z. Liu, C. Legrand, L. Fievez, F. Ginhoux, C. Radermecker, F. Bureau, and T. Marichal. 2023. MafB-restricted local monocyte proliferation precedes lung interstitial macrophage differentiation. Nat Immunol. 24:827–840. doi:10.1038/s41590-023-01468-3.

Wang, G.G., K.R. Calvo, M.P. Pasillas, D.B. Sykes, H. Häcker, and M.P. Kamps. 2006. Quantitative production of macrophages or neutrophils ex vivo using conditional Hoxb8. Nature Methods. 3:287–293. doi:10.1038/nmeth865.

Wang, X., S. Pesakhov, J.S. Harrison, M. Kafka, M. Danilenko, and G.P. Studzinski. 2015. The MAPK ERK5, but not ERK1/2, inhibits the progression of monocytic phenotype to the functioning macrophage. Experimental Cell Research. 330:199–211. doi:10.1016/j.yexcr.2014.10.003.

Yang, S., J. Zeng, W. Hao, R. Sun, Y. Tuo, L. Tan, H. Zhang, R. Liu, and H. Bai. 2023. IL-21/IL- 21R Promotes the Pro-Inflammatory Effects of Macrophages during C. muridarum Respiratory Infection. Int J Mol Sci. 24:12557. doi:10.3390/ijms241612557.

Zhang, G., N.A. deWeerd, S.A. Stifter, L. Liu, B. Zhou, W. Wang, Y. Zhou, B. Ying, X. Hu, A.Y. Matthews, M. Ellis, J.A. Triccas, P.J. Hertzog, W.J. Britton, X. Chen, and C.G. Feng. 2018. A proline deletion in IFNAR1 impairs IFN-signaling and underlies increased resistance to tuberculosis in humans. Nat Commun. 9:85. doi:10.1038/s41467-017-02611-z.

Zhou, Y., B. Zhou, L. Pache, M. Chang, A.H. Khodabakhshi, O. Tanaseichuk, C. Benner, and S.K. Chanda. 2019. Metascape provides a biologist-oriented resource for the analysis of systems-level datasets. Nat Commun. 10:1523. doi:10.1038/s41467-019-09234-6.

